# Dopamine release in nucleus accumbens core during social behaviors in mice

**DOI:** 10.1101/2021.06.22.449478

**Authors:** Bing Dai, Fangmiao Sun, Amy Kuang, Yulong Li, Dayu Lin

## Abstract

Social behaviors are among the most important and rewarding motivational behaviors. How dopamine, a “reward” signal, releases in the nucleus accumbens (NAc) during social behaviors has become a topic of interest for decades. However, limitations in early recording methods, such as microdialysis, prevented a complete understanding of moment-to-moment dopamine responses during social behaviors. Here, we employ a genetically encoded dopamine sensor, GRAB_DA2h_, to record dopamine activity in the NAc core in mice and find acute changes in extracellular dopamine levels during all three phases of social behaviors: approach, investigation and consummation. Dopamine release during approach phase correlates with animal’s motivation towards the conspecific whereas its release during consummatory phase signals the valence of the experience. Furthermore, dopamine release during sexual and aggressive behaviors shows sex differences that correlate with the potential value of those experiences. Overall, our results reveal rich and temporally precise motivation and value information encoded by NAc dopamine during social behaviors and beyond.

## Introduction

The dopaminergic input from ventral tegmental area (VTA) to nucleus accumbens (NAc) is well known for its relevance to reward, but how so? One function of NAc dopamine release is to signal reward expectation which could be used as a motivating signal. In support of this hypothesis, altering the dopamine release artificially increases the willingness to work (Hamid et al., 2016; Phillips et al., 2003). The other function of NAc dopamine is to signal errors in reward prediction, providing a learning signal to guide future behavior (Bayer and Glimcher, 2005; Eshel et al., 2016). In cases when a reward arrives unexpectedly or in other words, the predicted value is zero, the reward prediction error is equivalent to the actual value of a stimulus. Thus, dopamine could signal the motivation to achieve a goal before the goal is obtained and the value of the goal upon obtaining it.

Social behaviors, such as sexual and parental behaviors, are among the most important and rewarding motivated behaviors (Trezza et al., 2011). The end goals of these behaviors, e.g., reproduction and fostering youngsters, are essential for the survival of a species. Thus, animals are innately motivated to engage in social behaviors and in some cases, willing to work hard for such opportunities. Social behaviors are intrinsically rewarding and can serve as unconditioned stimulus (US) for associative learning. For example, male and female rodents will establish a preference for the context associated with copulation and postpartum female rats will learn to lever press to gain access to a pup (Hauser and Gandelman, 1985; Tzschentke, 2007; Wilsoncroft, 1968).

Many studies have been carried out since the early 90s to ask whether and how dopamine level changes in NAc during social behaviors. Early microdialysis studies with a temporal resolution of minutes revealed a gradual increase of dopamine in NAc in male rats that copulated to ejaculation with receptive females (Damsma et al., 1992; Fumero et al., 1994; Pfaus et al., 1990; Pleim et al., 1990; Wang et al., 1995; Wenkstern et al., 1993). Later voltammetry recording with a higher temporal resolution (subsecond to second) found that dopamine transients increase mainly during initial female encounter but not during consummatory phase of sexual behaviors, such as deep thrust (Robinson et al., 2002). This is surprising given that consummatory sexual actions are required for establishing conditioned place preference – typically a dopamine-dependent learning process (Kippin and Pfaus, 2001; Tenk et al., 2009). Recently, we used a genetically encoded dopamine sensor, namely GRAB_DA_, to optically record the dopamine signal in NAc in male mice with millisecond resolution and found time-locked dopamine increase during each episode of thrust and ejaculation, supporting a role of dopamine in encoding the hedonic value of sexual behaviors (Sun et al., 2018; Sun et al., 2020). This result suggests that the early recording methods may lack the temporal resolution and sensitivity to reveal the full details of dopamine responses during social behaviors.

Similar to male sexual behaviors, dopamine increase in NAc during female sexual behaviors and pup interactions have been reported using microdialysis and to a lesser extent voltammetry (Afonso et al., 2008; Afonso et al., 2009; Afonso et al., 2013; Becker et al., 2001; Champagne et al., 2004; Hansen et al., 1993; Jenkins and Becker, 2003; Kohlert and Meisel, 1999; Lavi-Avnon et al., 2008; Meisel et al., 1993; Mermelstein and Becker, 1995; Pfaus et al., 1995; Shnitko et al., 2017). These studies unequivocally suggested that the dopamine release in NAc is correlated with the animal’s sexual or maternal motivation (Afonso et al., 2009; Champagne et al., 2004; Kohlert and Meisel, 1999; Mermelstein and Becker, 1995). However, considering the poor temporal resolution of microdialysis, dopamine responses during individual behavioral events, especially those lasting for just a second or two, remain unclear. It also remains to be determined whether the slow changes in dopamine levels truly reflects slow dynamics of dopamine or whether they simply reflect methodological limitations. Furthermore, nearly all studies on parental behaviors have focused on mothers probably due to the fact that mother is the main care giver. One study showed that dopamine release to pups in NAc in naïve and pair-bonded male prairie voles are quantitatively similar although pair-bonded males show enhanced paternal behavior (Lei et al., 2017). These results raise the question of whether pup-triggered dopamine release in males varies with the paternal state as is the case in females (Afonso et al., 2008; Afonso et al., 2009; Champagne et al., 2004).

Aggression is another important type of motivational behaviors towards a social target. Animals are willing to work for the opportunity to attack a conspecific, especially when the outcome of attack is likely winning (Falkner et al., 2016; Fish et al., 2005; Fish et al., 2002; Fish et al., 2008; Golden et al., 2017; Golden et al., 2019c; May and Kennedy, 2009). Winning also supports associative learning: animals demonstrate a preference for the context where winning occurs (Aleyasin et al., 2018; Golden et al., 2016; Golden et al., 2019c; Martinez et al., 1995; Stagkourakis et al., 2018). Several studies investigated changes in dopamine levels related to aggressive behaviors and found a slow and sustained increase in NAc (Beiderbeck et al., 2012; van Erp and Miczek, 2000). However, unlike sexual and parental behaviors, dopamine rises slowly and can remain elevated for over an hour (Beiderbeck et al., 2012; van Erp and Miczek, 2000). Whether the particularly slow dynamics reflect a qualitative difference in dopamine release associated with aggressive vs. other social behaviors or whether they are merely caused by low sampling rates remain to be investigated. Further complicating the findings is the observation that NAc dopamine also increases after defeat (Tidey and Miczek, 1996). Since defeat is clearly a negative experience, it suggests that dopamine increase during social behaviors may signal salience instead of valence. Indeed, dopamine in NAc may signal both salience and valence depending on the subregion. While dopamine increases to stimuli of both positive and negative valence in the ventral NAc shell, dopamine only signals positive valence in other parts of NAc (de Jong et al., 2019; Yuan et al., 2019).

Taken together, while dopamine release in NAc during social behaviors has been a topic of interest for the last three decades, many questions remain unaddressed largely due to technical limitations, heterogeneity of release pattern in NAc and differences in methodological details across studies. Thus, the goal of our current study is to comprehensively investigate the dopamine release in NAc during social behaviors using an optical recording method with fast temporal resolution and cell type specificity. By recording from both males and females, our study also examined potential sexual dimorphism in dopamine release during social behaviors. Here, we specifically focused on NAc core given that this region is known to signal both motivation and reward, two important variables relevant for social behaviors (Hamid et al., 2016).

## Results

### Dopamine release in NAc core during approach and investigation

To engage in social behaviors, animals need to first reach a social target. In laboratory settings, this “approach” phase mainly consists of one animal walking towards another animal in the same arena although more complicated movements, e.g., lever pressing or grid cross, can also be involved (Golden et al., 2019c; Trezza et al., 2011; Wei et al., 2021). The seeking behavior, most common approaching behavior, reflects an animal’s motivation to a social target and the relevant consummatory social actions. Upon gaining access to the social target, the animal investigates it closely. After variable time of investigation, the consummatory action is initiated. Although interaction with a social target alone can be rewarding, the consummatory social actions are of higher hedonic value as social behavior-dependent associative learning typically requires the successful completion of consummatory social actions (Trezza et al., 2011).

To understand how dopamine levels change during different stages of social behaviors, we performed optical recording of the dopamine signal by virally expressing Cre-dependent (Cre-on) GRAB_DA2h_, a genetically encoded fluorescent DA sensor, unilaterally or bilaterally in the NAc core of Drd1-Cre mice, and implanting 400-μm multimode optic fiber(s) immediately above the virus injection site(s) for light delivery and collection (Sun et al., 2018; Sun et al., 2020). In a subset of animals, we also injected Cre-on or Cre-off GRAB_DA2h_ virus into the contralateral side of the NAc core to compare dopamine release onto D1R and non-D1R cells (**Figure 1a-1c**). Control animals were injected with Cre-on GFP virus on both sides. Only animals with correct fiber targeting are included in the analysis (**Figure 1 – figure supplement 1**).

**Figure 1.**
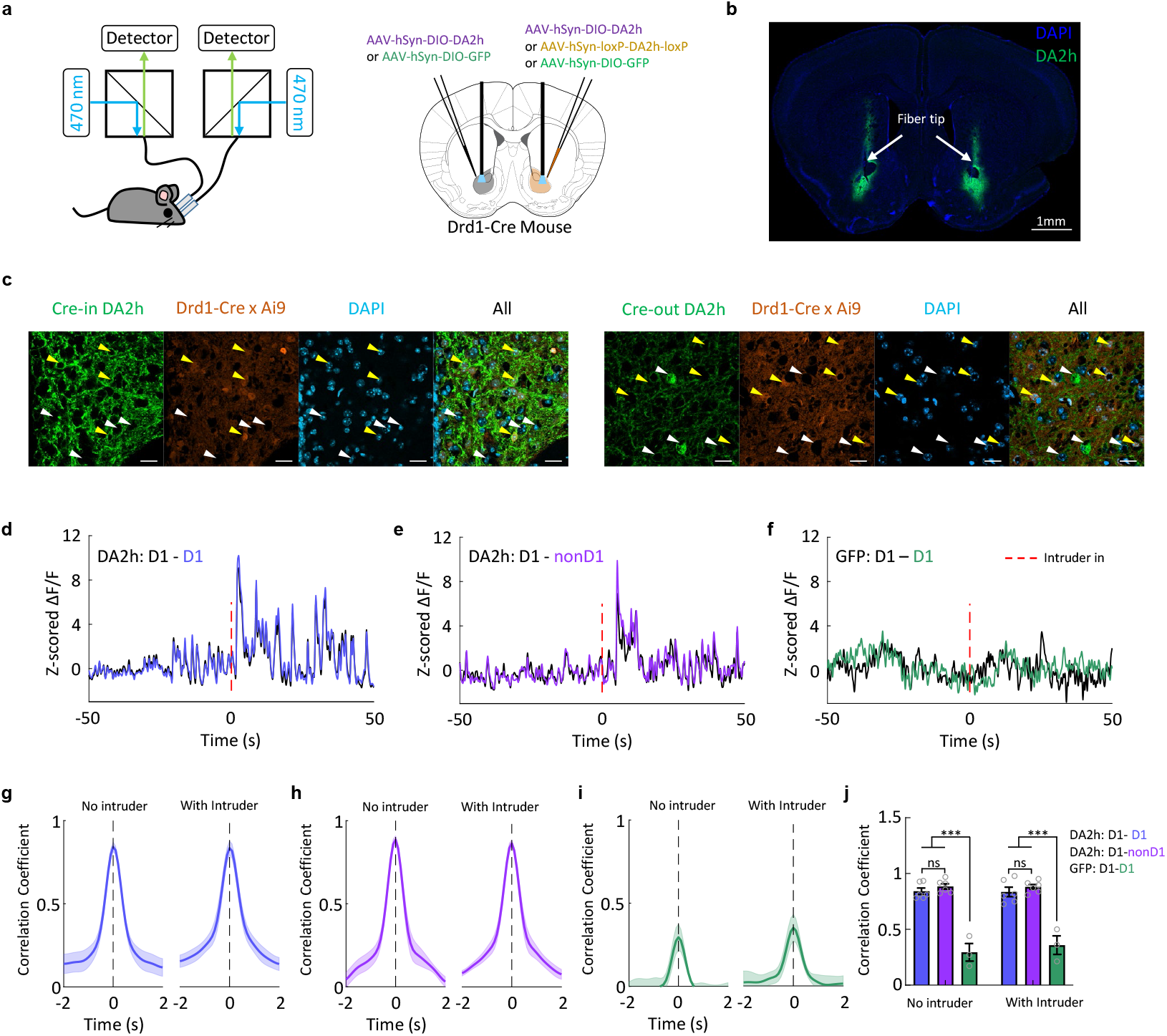
DA release sensed by D1 cells and non-D1 cells are highly correlated. **a**, Schematic illustration of the recording setup and virus injection. Atlas image is adopted from (Franklin and Paxinos, 2013) **b**, Representative image showing the expression of GRAB_DA2h_ in both hemispheres and optic fiber tracks. (Scale bar, 1mm.) **c**, Representative images showing the expression of GRAB_DA2h_ in D1 cells (left, yellow arrows) in an animal injected with Cre-in virus and in non-D1 cells (right, white arrows) in an animal injected with Cre-out virus. (Scale bar, 20μm.) **d**, Representative traces of GRAB_DA2h_ from D1R cells in both hemispheres. Time 0 indicates when a conspecific intruder is introduced. Similar results were observed from six mice. **e**, Representative traces of GRAB_DA2h_ from D1R cells in one hemisphere and that of non-D1R cells in the other hemisphere. Time 0 indicates when a conspecific intruder is introduced. Similar results were observed from six mice. **f**, Representative traces of GFP from D1R cells in both hemispheres. Similar results were observed from three mice. **g**, The time shifted correlation coefficient between the GRAB_DA2h_ signals from two hemispheres before and after the introduction of an intruder. Shaded area: s.e.m. (n=6 mice.) **h**, The time shifted correlation coefficient between GRAB_DA2h_ in D1R and non-D1R cells in two different hemispheres before and after an intruder introduction. Shaded area: s.e.m. (n=6 mice.) **i**, The time shifted correlation coefficient between GFP in D1R cells from different hemispheres before and after intruder introduction. Shaded area: s.e.m. (n=3 mice.) **j**, Group summary of peak correlation coefficients of GRAB_DA2h_ or GFP signals between hemispheres. Mean ± s.e.m overlaid for each group. (n= 6 mice for each GRAB_DA2h_ group and n=3 for the GFP group. Bonferroni’s multiple comparisons following two-way ANOVA. ***p<0.001.)

In animals with bilateral GRAB_DA_ expression in D1R cells of NAc core, we observed highly correlated activity regardless of the presence of an intruder animal, suggesting synchronized dopamine release in the two hemispheres (**Figure 1d, 1g and 1j**). Furthermore, in animals with GRAB_DA_ expression in D1R and non-D1R cells in the contralateral sides of NAc core, we observed similarly highly correlated dopamine signals, suggesting that D1R and non-D1R cells are likely sense similar level of dopamine fluctuation (**Figure 1e, 1h and 1j**). Of note, this does not rule out the possibility that D1R and non-D1R cells sense different dopamine inputs at the microscopic level since it is beyond the spatial resolution allowed by our recording method (Liu et al., 2018). The highly synchronized activity was not due to signal fluctuation related to locomotion. In animals with bilateral GFP expression, we observed significantly lower correlation in fluorescence signals between the two hemispheres (**Figure 1f, 1i and 1j**). Given the highly correlated dopamine signal sensed by D1R and non-D1R cells, our subsequent recordings were only obtained from D1R cells in NAc core (**Figures 2a-2c**).

**Figure 2.**
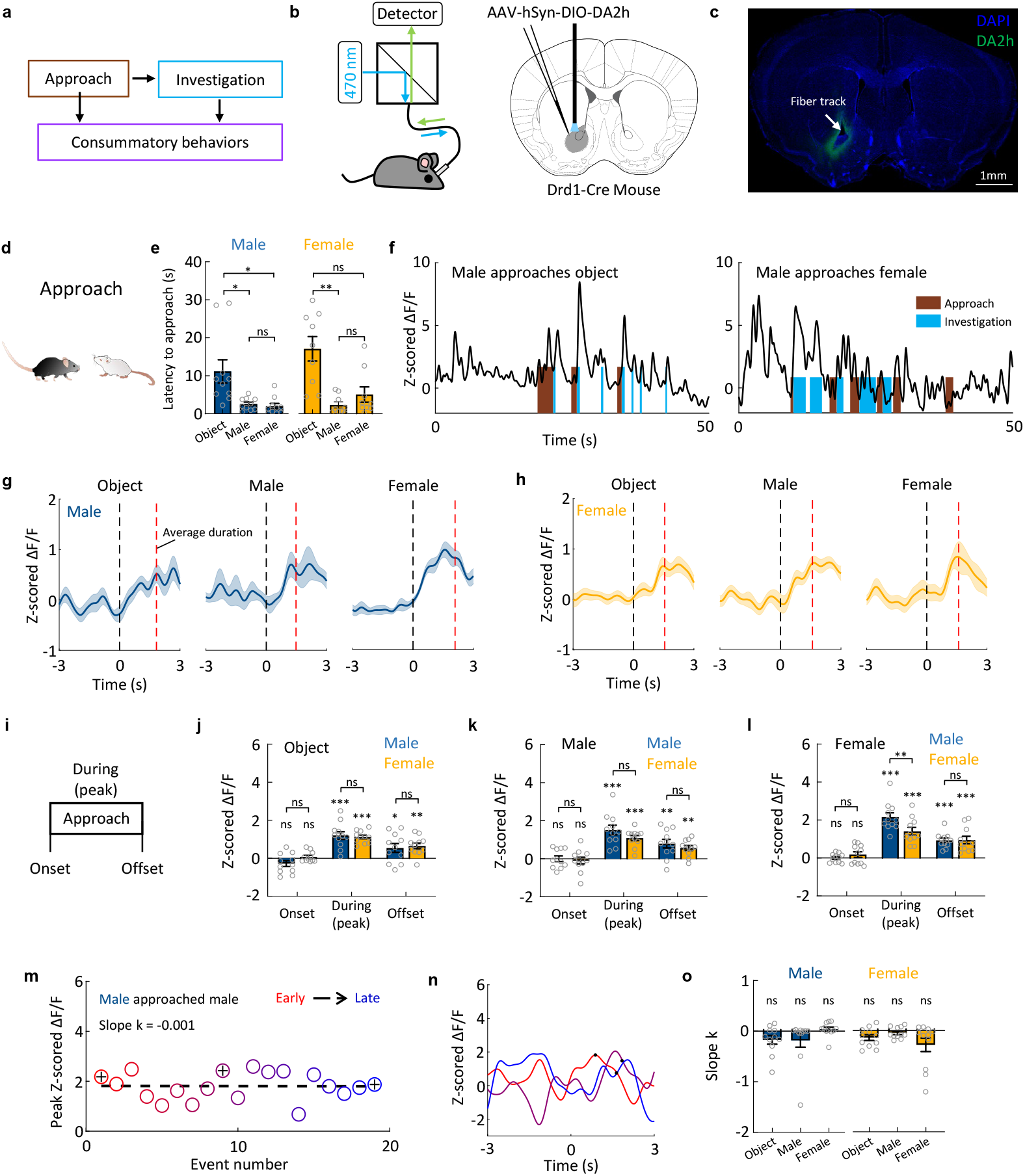
DA responses during approaching social and non-social targets. **a**, Schematics illustrating different stages of social behaviors. **b**, Schematics illustrating the experimental design. **c**, Representative image showing the expression of GRAB_DA2h_ and optic fiber track. (Scale bar, 1mm.) **d**, A cartoon illustration of approaching behavior. **e**, Group summary of the latency to approach towards different targets in male (blue) and female mice (yellow). Mean ± s.e.m overlaid with individual data points for each group. (n=10 male mice, Friedman test, p=0.007; n=10 female mice, Friedman test, p=0.006. Dunn’s multiple comparisons post-tests revealed the difference within male and female groups, *p<0.05, **p<0.01) **f**, Representative traces of Z scored ΔF/F of GRAB_DA2h_ during social interaction from an example male mouse. Color shades indicate annotated behaviors. Similar results were observed from 11 male mice and 11 female mice. **g**, Average post-event histograms aligned to the onset of approach for male mice. Shaded area: s.e.m. (n=11 mice.) **h**, Average post-event histograms aligned to the onset of approach for female mice. Shaded area: s.e.m. (n=11 mice.) **i**, Schematics showing the time periods used for characterizing DA responses related to approach. **j-l**, Summary of Z scored GRAB_DA2h_ responses at the onset and offset of, and during approach a novel object (j), a male conspecific (k) and a female conspecific (l). (n=11 male mice and n=11 female mice. One sample t test followed by FDR correction (FDR = 0.05) to reveal significant responses. *p<0.05, **p<0.01, ***p<0.001. Bonferroni’s multiple comparisons following two way ANOVA revealed no difference in responses between males and females. **p<0.01) **m**, A scatter plot showing peak response during repeated approach trials from one example male-male encounter session. **n**, Representative GRAB_DA2h_ traces from early and late approach bouts. Black dots indicate the offset of behavior episodes. **o**, Summary of slope k of GRAB_DA2h_ responses over repeated approach episodes. (n=11 male mice and n=11 female mice. One sample Wilcoxon test followed by FDR correction (FDR = 0.05).

During each recording session, we sequentially introduced a conspecific male, a female, and a novel object into the tested mouse’s home cage, each for 10 minutes or until ejaculation was achieved in the case of opposite sex interaction. Upon introduction of a stimulus, the home-cage animal approached the target quickly. For both male and female, the latency to approach a social target was shorter than that towards an object (**Figures 2d-2e**). At the approach onset -- defined as the first step towards the target, the dopamine level was not elevated, suggesting that dopamine increase was unlikely to play a role in initiating approach (**Figure 2f-2l**). During approach, dopamine levels gradually increased (**Figure 2g-2h**). For males, the maximum dopamine increase during approaching a female was significantly higher than that during approaching an object (Female vs Object: p = 0.04, one-way ANOVA followed by Tukey’s multiple comparisons test) whereas females showed comparable dopamine increase during approach towards all targets (**Figure 2j-2l**). At the offset of approach, which is often followed by other social behaviors, e.g., investigation, the dopamine remained significantly elevated (**Figure 2j-2l**). We next asked whether the dopamine increase during approach adapted over trials and found no consistent decrease especially towards a social target of the opposite sex (**Figure 2m-o**).

To address whether the response during approach was due to movement per se, we tracked the position of the test animal in the absence of an intruder and identified time points when the animal initiated locomotion (**Figure 2 – figure supplement 1a-1b**). No increase in dopamine activity was observed at the onset of locomotion (**Figure 2 – figure supplement 1c-1d)** In fact, dopamine slightly but significantly decreased when the animal initiated locomotion and the movement velocity was significantly negatively correlated with the dopamine signal, suggesting that the dopamine increase during approach was not due to locomotion itself (**Figure 2 – figure supplement 1c-1f**).

Upon reaching the target, test animals closely investigate the stimulus. For both females and males, the average duration of investigation bout was longer towards social targets than that towards novel objects (**Figure 3b**). Dopamine levels at the onset of investigation were already significantly elevated above the baseline, likely due to the dopamine increase during approach (**Figures 3c-3m**). During investigation, the dopamine further rose transiently before it dropped (**Figures 3d-3m**). At the offset of social investigation, the dopamine level largely returned to the baseline level in males and is slightly elevated in females (**Figure 3k-3m**). The peak dopamine increases during investigation of social targets were significantly higher than that during object investigation (Male: male vs object: p<0.001; female vs object: p = 0.009. Female: male vs object: p = 0.01; female vs object: p = 0.005. One-way ANOVA followed by Tukey’s multiple comparisons test). Dopamine response during investigation adapted quickly regardless of the targets (**Figures 3n-3p**). On average, dopamine increase during the tenth investigation bout was around 20-40% of the first bout (**Figure 3q**).

**Figure 3.**
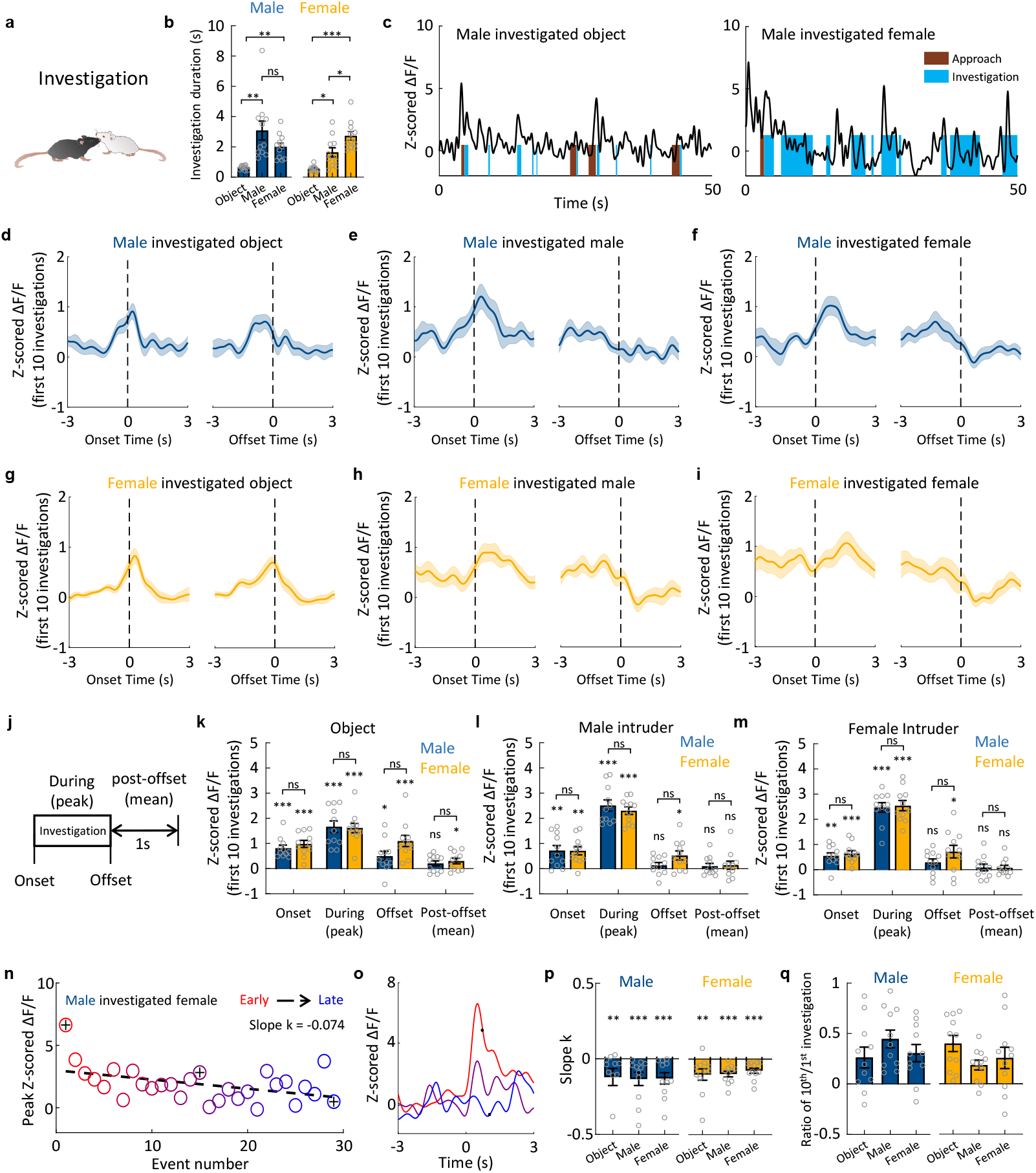
DA responses during male and female investigatory behaviors. **a**, A cartoon illustration of social investigation. **b**, Summary of average investigation durations towards various targets in male and female mice. Mean ± s.e.m overlaid with individual data points for each group. (n=11 male mice, one-way ANOVA, F(1.203, 12.03)=9.151, p=0.008; n=11 female mice, one-way ANOVA, F(1.938,19.38)=24.43, p<0.001. Tukey’s multiple comparisons post-tests revealed the differences within male and female groups, *p<0.05, **p<0.01, ***p<0.001.) **c**, Representative traces of Z scored ΔF/F of GRAB_DA2h_ during object and female interaction from an example male mouse. Similar results were observed from 11 male mice and 11 female mice. **d-i**, Average post-event histograms aligned to the onset (left) and offset (right) of investigation towards object (**d**), a male intruder (**e**) and a female intruder (**f**). Shaded area: s.e.m. (n=11 male mice and 11 female mice.) **j**, Schematics showing the time periods used for characterizing DA responses related to investigation. **k-m**, Summary of Z scored GRAB_DA2h_ responses at the onset and offset of, during and after investigating a novel object (**k**), a male mouse (**k**) and a female mouse (**m**). Signal changes across bouts. Mean ± s.e.m overlaid with individual data points for each group. (n=11 male mice and n=11 female mice. One sample t test followed by FDR correction revealed significant responses, *p<0.05, **p<0.01, ***p<0.001. Bonferroni’s multiple comparisons following two-way ANOVA revealed no difference between male’s and female’s responses.) **n**, A scatter plot showing peak responses during repeated investigation trials from one example session of male-female encounter. **o**, Representative GRAB_DA2h_ traces from early and late investigation bouts. Black dots mark the end of investigation. **p**. Summary of slope k of GRAB_DA2h_ responses over repeated investigation events. Mean ± s.e.m overlaid with individual data points for each group. (n=11 male mice and n=11 female mice. One sample Wilcoxon rank test followed by FDR correction revealed significant adaptations. **p<0.01, ***p<0.001.) **q**, The ratio of peak DA signals during the tenth investigation bout to that of the first bout. Mean ± s.e.m overlaid individual data points for each group. (n=11 male mice and n=11 female mice.)

### Dopamine release in NAc core during sexual behaviors

During male and female encounters, after a period of investigation, male mice initiate mounting towards the females. We used sexually experienced males and they often attempt to mount females regardless of the female’s receptivity level. In these males, the dopamine dynamic during mounting towards receptive and non-receptive females was similar: it was elevated at the mounting onset, continued to rise transiently during mounting before dropping below the baseline at the offset of mounting if the mounting did not transition into intromission (**Figures 4a-4e and 4j-4l**). Furthermore, the dopamine suppression remained for at least one second after mounting termination (**Figures 4k-4l**). A different pattern of dopamine release was observed in females. Both non-receptive and receptive females showed no change in dopamine level at the onset of being mounted (**Figures 4d-4e and 4k-4l**) When being mounted, non-receptive female showed a decrease in dopamine which was maintained throughout the behavior and for at least one second after male stopped mounting (**Figures 4c-4d and 4k**). In contrast, when being mounted, receptive females showed a transient increase in dopamine which returned to baseline at the offset (**Figures 4c, 4e and 4l**)

**Figure 4.**
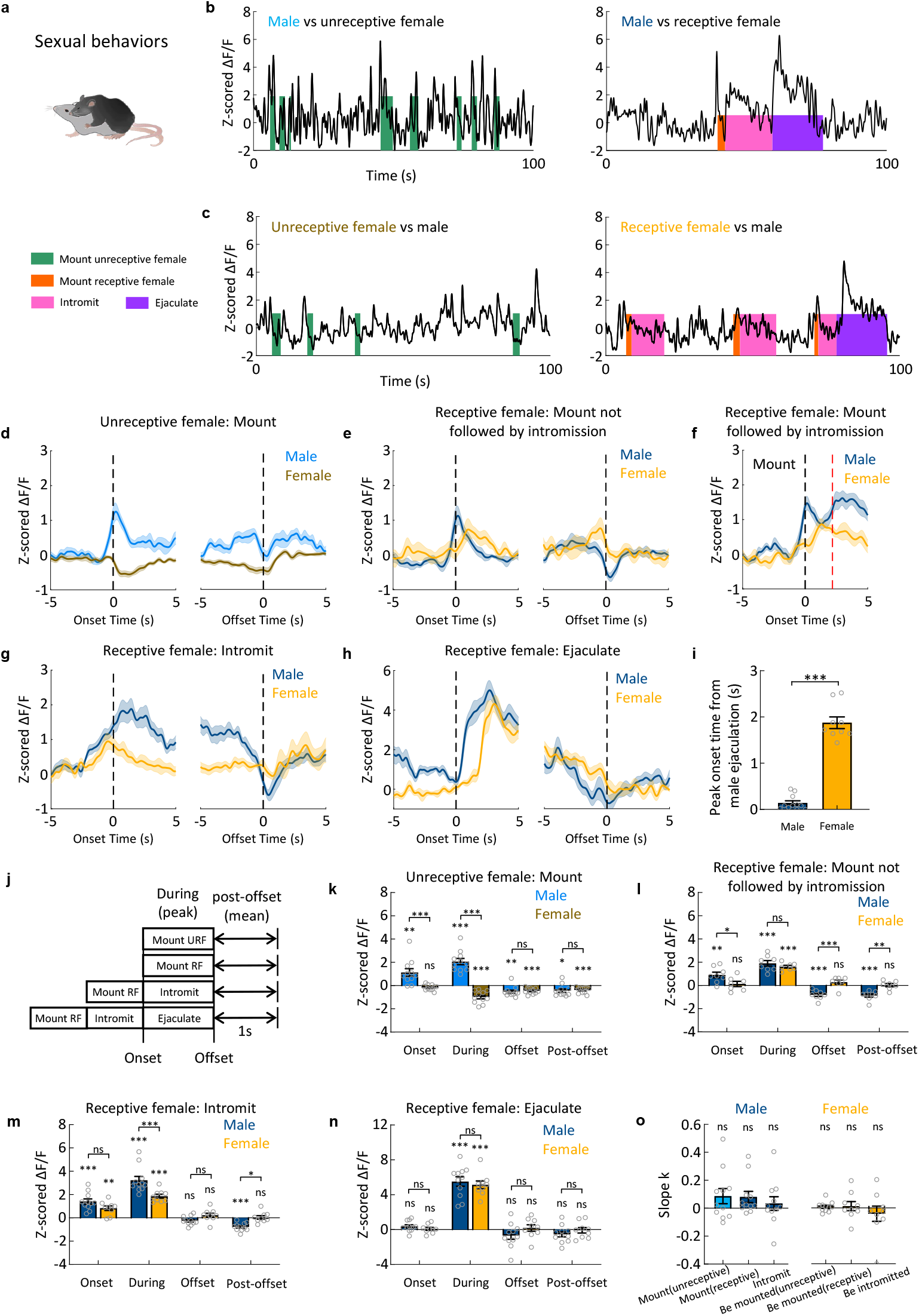
DA responses during male and female sexual behaviors. **a**, A cartoon illustration of mouse mating. **b-c**, Representative traces of Z scored ΔF/F of GRAB_DA2h_ during various stages of mating in male and female mice. Similar results were observed from 11 male mice (**b**), 11 unreceptive female mice(**c, left**), and 9 receptive female mice (**c, right**). **d**, Average post-event histograms aligned to the onset (left) and offset (right) of the male mounting events, when the female mice were unreceptive. Shaded area: s.e.m. (n=11 male mice and n=11 female mice.) **e-h**, Average post-event histograms aligned to the onset (left) and offset (right) of the indicated mating events, when the female mice were receptive. Shaded area: s.e.m. (n=11 male mice and n=9 female mice.) **i**, Bar graphs showing the latency from the onset of ejaculation to the moment when DA signals start to increase. Mean ± s.e.m overlaid with individual data points for each group. (n=11 male mice and n=9 female mice, Mann-Whitney test, ***p<0.001) **j**, Schematics showing the time periods used for characterizing DA responses associated with mating. **k**. Summary of Z scored GRAB_DA2h_ responses at the onset and offset of, during and after male mounting when the females were non-receptive. Mean ± s.e.m overlaid with individual data points for each group. (n=11 male mice and n=11 female mice. One sample t test followed by FDR correction to reveal significant responses, *p<0.05, **p<0.01, ***p<0.001. Bonferroni’s multiple comparisons following two-way ANOVA revealed differences in responses between males and females, ***p<0.001.) **l**, Summary of Z scored GRAB_DA2h_ responses at the onset and offset of, during and after male mounting that was not followed by intromission. The female mice were receptive. Mean ± s.e.m overlaid with individual data points for each group. (n=8 male mice and n=7 female mice. One sample t test followed by FDR correction to reveal significant responses, *p<0.05, **p<0.01, ***p<0.001. Bonferroni’s multiple comparisons following two-way ANOVA revealed differences in responses between males and females, *p<0.05, **p<0.01, ***p<0.001.) **m**, Summary of Z scored GRAB_DA2h_ responses at the onset and offset of, during and after male intromission that was not followed by ejaculation. The female mice were receptive. Mean ± s.e.m overlaid with individual data points for each group. (n=11 male mice and n=9 female mice. One sample t test followed by FDR correction to reveal significant responses, **p<0.01, ***p<0.001. Bonferroni’s multiple comparisons following two-way ANOVA revealed differences in responses between males and females, *p<0.05, ***p<0.001.) **n**, Summary of Z scored GRAB_DA2h_ responses at the onset and offset of, during and after male ejaculation. Mean ± s.e.m overlaid with individual data points for each group. (n=11 male mice and n=9 female mice. One sample t test followed by FDR correction to reveal significant responses, ***p<0.001. Bonferroni’s multiple comparisons following two-way ANOVA revealed no difference in responses between males and females.) **o**, Summary of slope k of GRAB_DA2h_ responses over repeated mating events. Mean ± s.e.m overlaid with individual data points for each group. (n=11 male mice, n=11 unreceptive female mice, and n=9 receptive female mice. One sample t test followed by FDR correction revealed no significant adaptation in response magnitude over trials.)

If the females are receptive, males advance mounting to intromission, a rhythmic pelvic movement presumably resulting in penile insertion. At the onset of intromission (the same moment as mounting offset), dopamine level was significantly elevated and continued to rise for approximately one second before it gradually decreased to the baseline level at the offset of intromission (**Figures 4b, 4f-4g and 4m**). If the intromission did not transition to ejaculation, dopamine decreased below the baseline for at least one second after intromission terminated (**Figures 4g and 4m**). In females, while the dopamine level was also elevated at the onset of being intromitted, it gradually and monotonically decreased and returned to the baseline at the offset of being intromitted without further post-behavior change (**Figures 4c, 4g and 4m**). After repeated intromission, males achieve ejaculation which is characterized as a sudden cessation of all movements. Ejaculation resulted in the highest dopamine release in both males and females, but there was a noticeable temporal difference (**Figures 4b-4c, 4h-4i and 4n**). While dopamine increased sharply in males at the onset of ejaculation, the dopamine increase in females occurred approximately 2 s after the male ejaculation (**Figure 4i**). Within 5 seconds after the onset of male ejaculation, dopamine reached its peak level and then gradually decreased (**Figure 4h**). At the ejaculation offset, defined as the moment when male resumes movement, dopamine levels have returned to baseline and no further post-ejaculation dopamine changes were observed (**Figure 4n**). Lastly, in contrast to the gradual decrease of dopamine release during investigation, the peak dopamine release during sexual actions was either unchanged or slightly increased over repeated trials (**Figure 4o**).

### Dopamine release in NAc core during aggressive behaviors

When the males encounter male intruders, they initiate attack after a period of investigation. At the onset of each attack, dopamine levels were already elevated, likely due to increase during approach or investigation (**Figures 5b-5c, 5f**). During attack, dopamine rose transiently, reached peak and then dropped (**Figures 5b-5c and 5f**). At the offset of attack, dopamine levels remained above the baseline but quickly returned to pre-attack levels within one second (**Figure 5b-5c and 5f**). The maximum dopamine increase during male aggression was quantitatively comparable to the dopamine increase during sexual intercourse and consuming palatable food (i.e. peanut butter), supporting the notion that aggression could be rewarding (**Figure 5- figure supplement 1**).

**Figure 5.**
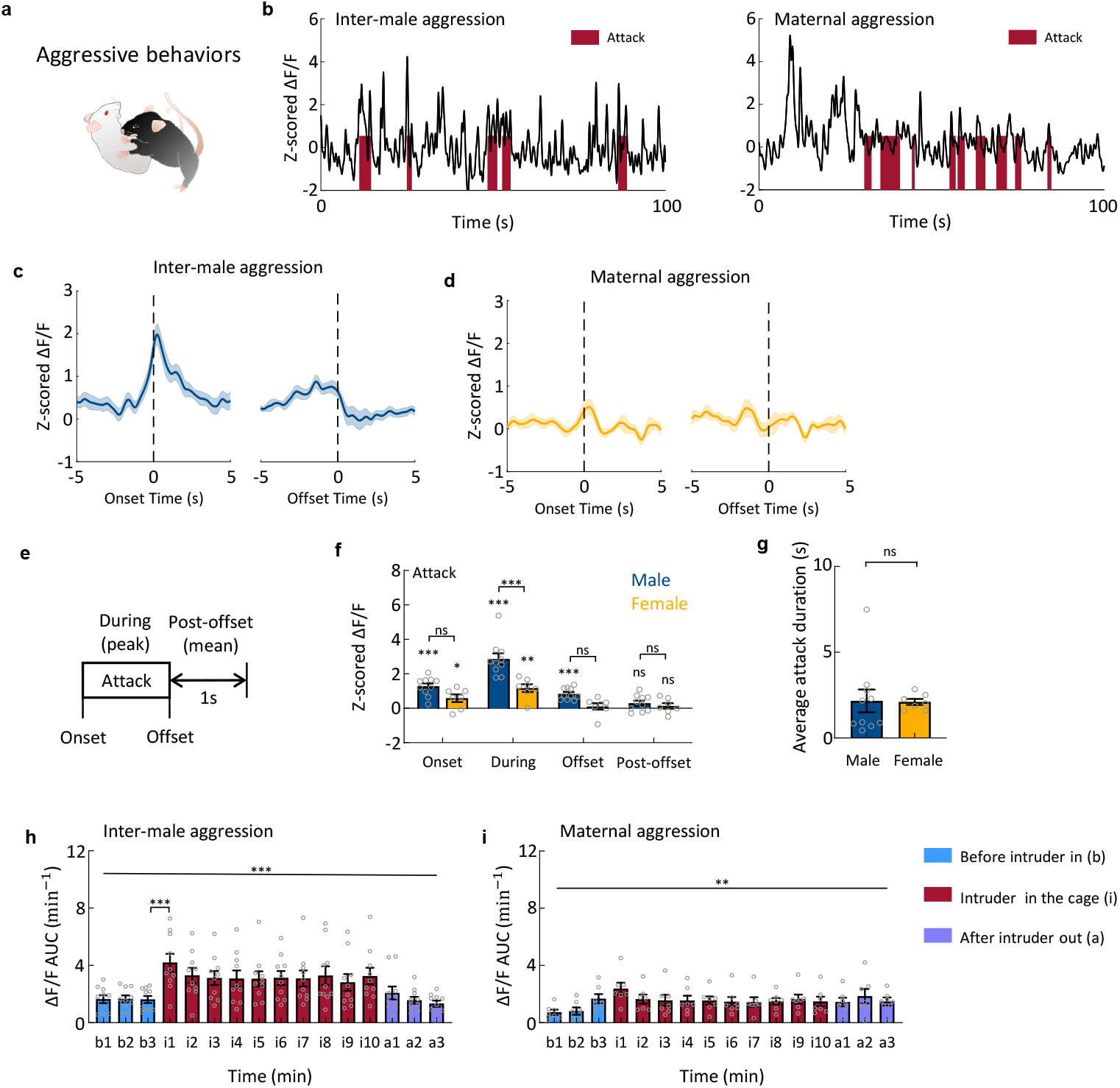
DA responses during aggressive behaviors. **a**, A cartoon illustration of mouse attack. **b**, Representative traces of Z scored ΔF/F of GRAB_DA2h_ during inter-male aggression and maternal aggression. Similar results were observed from 10 male mice and 7 female mice. **c-d**, Average post-event histograms aligned to the onset (left) and offset (right) of attack. Shaded area: s.e.m. (n=10 male mice and n=7 female mice.) **e**, Schematics showing the time periods used for characterizing DA responses related to attack. **f**, Summary of Z scored GRAB_DA2h_ responses at the onset and offset of, during and after attack. Mean ± s.e.m overlaid with individual data points for each group. (n=10 male mice and n=7 female mice. One sample t test followed by FDR revealed significant responses, *p<0.05, **p<0.01, ***p<0.001. Bonferroni’s multiple comparisons following two-way ANOVA revealed differences in responses between males and females, ***p<0.001.) **g**, Bar graph showing average attack durations for male and female mice. Mean ± s.e.m overlaid with individual data points for each group. (n=10 male mice and n=7 female mice, Mann-Whitney test, p=0.94.) **h**. Accumulated DA signals before, during, and after the male-male interaction. Mean ± s.e.m overlaid with individual data points for each group. (n = 11 mice, Friedman test, p<0.001. Dunn’s multiple comparisons revealed the significant difference between b3 and i1, ***p<0.001.) **i**, Accumulated DA signals before, during, and after the interaction between the lactating female with a female intruder. Mean ± s.e.m overlaid with individual data points for each group. (n = 9 mice, Friedman test, p=0.002. Dunn’s multiple comparisons revealed no difference between b3 and any the other time points.)

Non-lactating female mice typically show little aggression towards conspecific intruders. During lactation, however, females show a marked increase in aggression towards all intruders except pups, a phenomenon known as maternal aggression (St John and Corning, 1973). Maternal aggression contains both offensive and defensive attacks and its main purpose is to protect the young (Ferrari et al., 2000; Flannelly and Flannelly, 1985). In comparison to male aggression, peak dopamine increase during female attack was significantly lower (**Figures 5f**).

At the offset of female attack, dopamine levels returned to baseline (**Figures 5b, 5d and 5f**). The sex difference in dopamine release during attack was not due to difference in attack duration: in both males and females, each attack bout lasted approximately 2 seconds (**Figure 5g**).

Previous microdialysis measurements suggest a sustained elevation of dopamine levels after male-male confrontation (Beiderbeck et al., 2012; van Erp and Miczek, 2000). We thus analyzed the accumulated dopamine signal before, during and after the inter-male aggression and maternal aggression (**Figure 5h-5i**). While dopamine levels were significantly elevated during initial encounter with the intruder in males, we found no significant difference in dopamine levels between the pre-intruder and post-intruder periods (**Figure 5h**). To ensure that our recording method can detect a sustained increase in dopamine, we i. p. injected dopamine transporter (DAT) inhibitor (GBR12909 20 mg/kg) in a subset of animals. DAT mediates dopamine reuptake and its blockage is known to cause an elevation of extracellular dopamine concentration (Westerink et al., 1987). Ten minutes after injecting DAT inhibitor but not saline, the GRAB_DA2h_ signal showed a consistent and sustained upward shift, supporting that our method was capable of detecting a general increase in dopamine level (**Figure 5 – figure supplement 2**).

### Dopamine responses in NAc during pup-directed behaviors

We also examined the dopamine release during pup-directed behaviors. Pup-directed behaviors are unique in that they show qualitative differences based on the motivational state of the animals. Naïve male mice either ignore or attack pups while fathers readily care for the young (e.g., quickly retrieve a wandering pup back to the nest; (Perrigo et al., 1990)). Virgin female mice from laboratory stocks typically do not attack pups but many show avoidance (Mann et al., 1983). With repeated pup exposure, however, virgin females can stop avoiding pups and care for them, a process known as pup sensitization (Rosenblatt, 1967). Mothers are strongly motivated for pups. They not only care for pups for most of the day but are willing to trade other high value rewards for pups (Trezza et al., 2011). Thus, the varying motivation to pups across different reproductive stage makes pups a particular useful social stimulus for evaluating the neural representation of motivation.

During recording, we introduced 5 - 7 pups, one at time, into the home cage of the recorded mouse for a total of 10 minutes. Among the 10 recorded naïve males, 5 ignored the pups (non-hostile males, NHM) and 5 attacked and killed pups (hostile males, HM). 10 (5 non-hostile and 5 hostile males) of those males were then paired with a female and became a father (father males, FM). We then repeated the recording of the test males between postpartum day 2 and 5 of their cohoused females. During recording, all fathers showed paternal behaviors including pup retrieval.

The temporal dynamics of dopamine release during pup approach were similar among naïve hostile males, naïve non-hostile males, and fathers: dopamine was not elevated at the onset of approach and increased during approach and reached the maximum level towards the end of approach (**Figures 6a-6c, 6f and 6h**). However, there was a significant difference in peak dopamine release during pup approach among the three groups of males. Specifically, dopamine increase was the highest in fathers and the lowest in hostile males (**Figure 6k**). Upon reaching the pup, the males closely interacted with the pups, including investigating, licking, and grooming the pups. As pups were often occluded by the body of the male, we did not attempt to distinguish these behaviors. During close interaction, dopamine level transiently increased and then returned to the baseline in all recorded animals and the peak level did not differ among groups (**Figure 6d, 6g, 6i and 6l**). In hostile males, dopamine level also transiently increased during infanticide although the peak increase was slightly lower than that during retrieval in fathers (p = 0.05, unpaired t test) (**Figures 6e, 6j and 6m-6n**). Interestingly, we noticed a transient dopamine suppression below the baseline after fathers completed retrieval and disengaged with the pups (**Figures 6n**).

**Figure 6.**
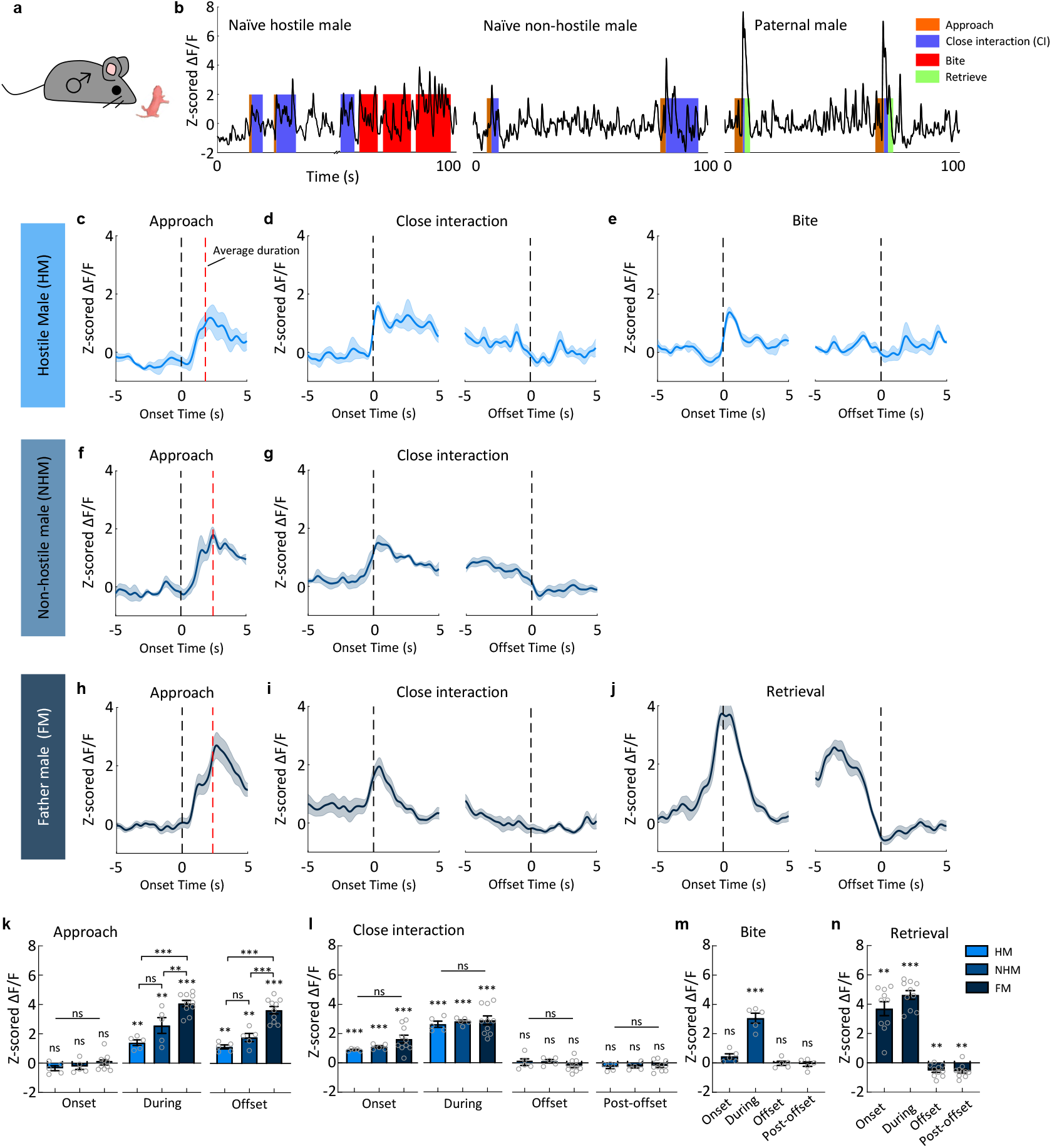
DA responses during pup-directed behaviors in naïve males and fathers. **a**, A cartoon illustration showing an adult male approaches a pup. **b**, Representative traces of Z scored ΔF/F of GRAB_DA2h_ during pup-directed behaviors in naive hostile males, naive non-hostile males, and fathers. Similar results were observed for 5 naive hostile male mice, 5 naive non-hostile male mice, and 10 father male mice. **c-e**, Average post-event histograms aligned to the onset (left) and offset (right) of the indicated pup-directed behaviors for the naive hostile male mice. Shaded area: s.e.m. (n=5 mice.) **f-g**, Average post-event histograms aligned to the onset (left) and offset (right) of the indicated pup-directed behaviors for the naive non-hostile male mice. Shaded area: s.e.m. (n=5 mice.) **h-j**, Average post-event histograms aligned to the onset (left) and offset (right) of the indicated pup-directed behaviors for fathers. Shaded area: s.e.m. (n=10 mice.) **k**, Summary of Z scored GRAB_DA2h_ responses at the onset and offset of, and during approach a pup for the naive hostile males, naive non-hostile males, and fathers. Mean ± s.e.m overlaid with individual data points for each group. (n=5 for naive hostile male mice, n=5 for naive non-hostile male mice, and n=10 for fathers. One sample t test followed by FDR correction to reveal responses, **p<0.01, ***p<0.001. Tukey’s multiple comparisons following one way ANOVA revealed the differences in responses between groups at the indicated moments, *p<0.05, **p<0.01, ***p<0.001.) **l**, Summary of Z scored GRAB_DA2h_ responses at the onset and offset of, during and after close interaction with a pup for the naive hostile male mice, naive non-hostile male mice, and fathers. Mean ± s.e.m overlaid with individual data points for each group. (n=5 for naive hostile male mice, n=5 for naive non-hostile male mice, and n=10 for father male mice. One sample t test followed by FDR correction to reveal significant responses, *p<0.05, **p<0.01, ***p<0.001. Tukey’s multiple comparisons following one-way ANOVA revealed no differences between groups at any moment. *p<0.05, **p<0.01, ***p<0.001.) **m**, Summary of Z scored GRAB_DA2h_ responses at the indicated moments of biting for the naive hostile male. Mean ± s.e.m overlaid with individual data points for each group. (n=5 male mice, one sample t test followed by FDR correction to reveal significant responses, ***p<0.001.) **n**, Summary of Z scored GRAB_DA2h_ responses at the indicated moments of retrieval for the fathers. Mean ± s.e.m overlaid with individual data points for each group. (n=10 male mice, one sample t test followed by FDR correction to reveal significant responses, *p<0.05, **p<0.01, ***p<0.001.)

Similarly, we examined dopamine release in NAc core in both naïve and lactating females (**Figure 7a**). 6/16 virgin females showed pup retrieval (naïve maternal females, NMF, latency to retrieve: 254.2 ± 108.5 s), while the rest of females only investigated and groomed the pups (naïve non-maternal females, NNF). All lactating females (10/10, mother females, MF) showed pup retrieval quickly after pup introduction (Latency to retrieve (mean ± STD): 23.51 ± 20.9 s). Dopamine release patterns during pup-interaction in females were qualitatively similar to that in males. Specifically, dopamine increased during pup approach, investigation, and retrieval **(Figure 7)**. The level of increase during pup approach was higher in females that showed maternal behaviors than those not (**Figure 7k**).

**Figure 7.**
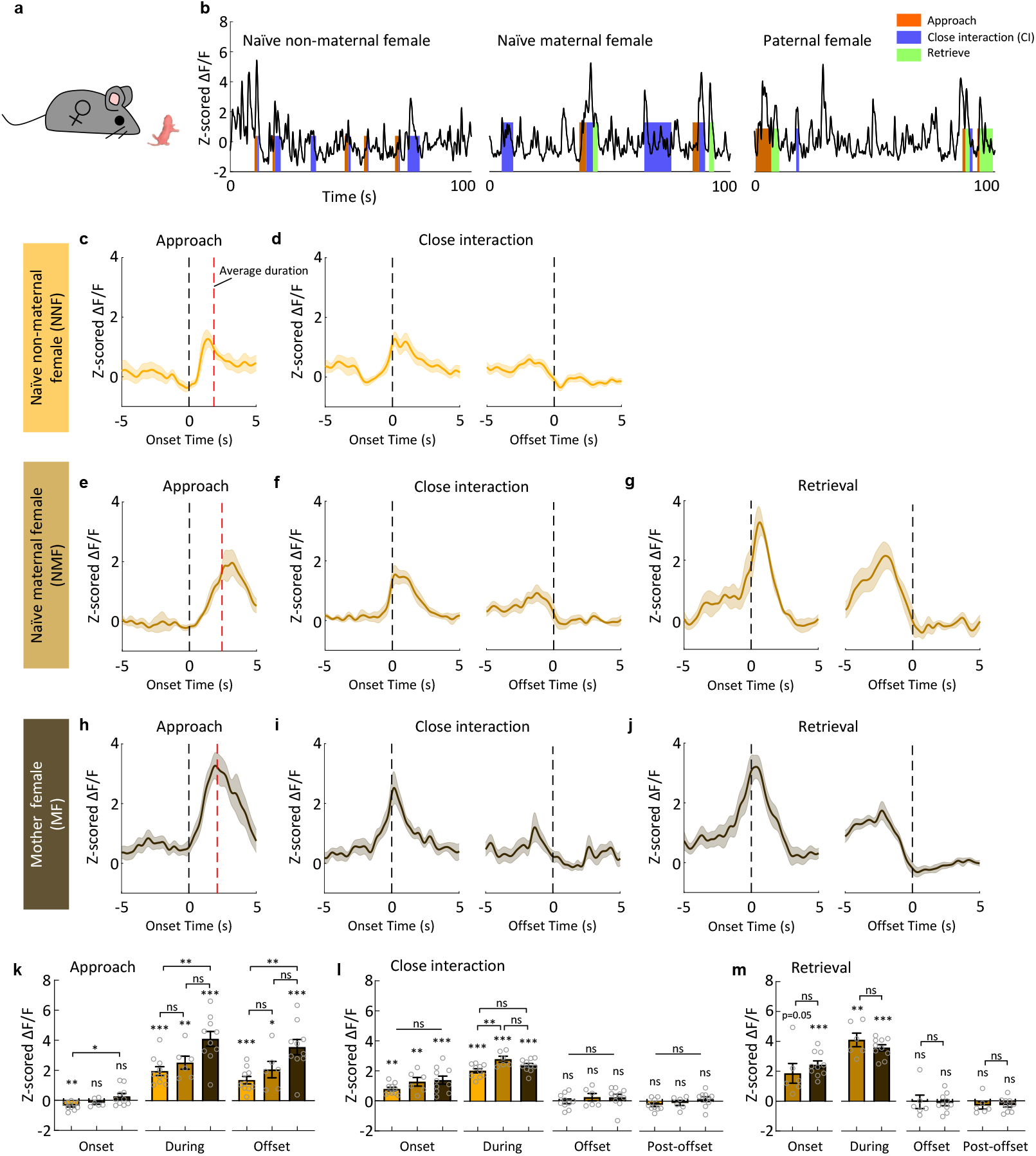
DA responses during pup-directed behaviors in naïve females and mothers. **a**, A cartoon illustration showing an adult female approaches a pup. **b**, Representative traces of Z scored ΔF/F of GRAB_DA2h_ during pup-directed behaviors in naive non-maternal female, naive maternal female, and paternal female mice. Similar results were observed from 10 naive non-maternal female mice, 6 naive maternal female mice, and 10 mother mice. **c-d**, Average post-event histograms aligned to the onset (left) and offset (right) of the indicated pup-directed events for the naive non-maternal female mice. Shaded area: s.e.m. (n=10 mice.) **e-g**, Average post-event histograms aligned to the onset (left) and offset (right) of the indicated pup-directed events for the naive maternal female mice. Shaded area: s.e.m. (n=6 mice.) **h-j**, Average post-event histograms aligned to the onset (left) and offset (right) of the indicated pup-directed events for mother mice. Shaded area: s.e.m. (n=10 mice.) **k**, Summary of Z scored GRAB_DA2h_ responses at the onset and offset of, and during approach a pup of the naive non-maternal female mice, naive maternal female mice, and paternal female mice. Mean ± s.e.m overlaid with individual data points for each group. (n=10 naive non-maternal female mice, n=6 naive maternal female mice, and n=10 mother female mice. One sample t test followed by FDR correction to reveal significant responses, *p<0.05, **p<0.01, ***p<0.001. Tukey’s multiple comparisons following one-way ANOVA revealed the differences between groups at the indicated moments, *p<0.05, **p<0.01, ***p<0.001.) **l**, Summary of Z scored GRAB_DA2h_ responses at the onset and offset of, during and after close interaction with a pup for the naive non-maternal female mice, naive maternal female mice, and paternal female mice. Mean ± s.e.m overlaid with individual data points for each group. (n=10 naive non-maternal female mice, n=6 naive maternal female mice, and n=10 mother female mice. One sample t test with FDR correction revealed significant responses, *p<0.05, **p<0.01, ***p<0.001. Tukey’s multiple comparisons following one-way ANOVA revealed the differences between groups at the indicated moments, **p<0.01.) **m**, Summary of Z scored GRAB_DA2h_ responses at the onset and offset of, during and after retrieving a pup for the naive maternal females and paternal females. Mean ± s.e.m overlaid with individual data points for each group. (n=6 naive maternal female mice, and n=10 mother female mice. One sample t test with FDR correction revealed significant responses,*p<0.05, **p<0.01, ***p<0.001. Unpaired t tests revealed no differences between groups of mice.)

Overall, these results suggest that dopamine increased acutely in NAc core during all phases of social behaviors. Dopamine increase during approach may signal the motivation towards a social target while dopamine increase during the consummatory social actions may signal the hedonic value of the behavior.

### Dopamine responses towards aversive experience

Do dopamine levels increase during all social behaviors or only the behaviors with positive valence? To address this question, we recorded GRAB_DA2h_ signal during defeat, a robust negative social experience. During recording, an aggressive Swiss Webster (SW) male mouse was introduced into the home cage of the recorded male for 10 minutes. Upon being attacked, the recording male mouse attempted to escape from the aggressor by flight and pushing and did not attempt to attack back. After a couple bouts of fighting, the recording male was clearly defeated: the SW intruder initiated all the attacks and the recorded mice stayed in corners and showed submissive postures. In contrast to the dopamine increase during attack, dopamine consistently and transiently decreased during defeat (**Figure 8b-d**). Interestingly, we noticed a rebound increase of dopamine after the termination of each defeat episode (**Figure 8b-d**).

**Figure 8.**
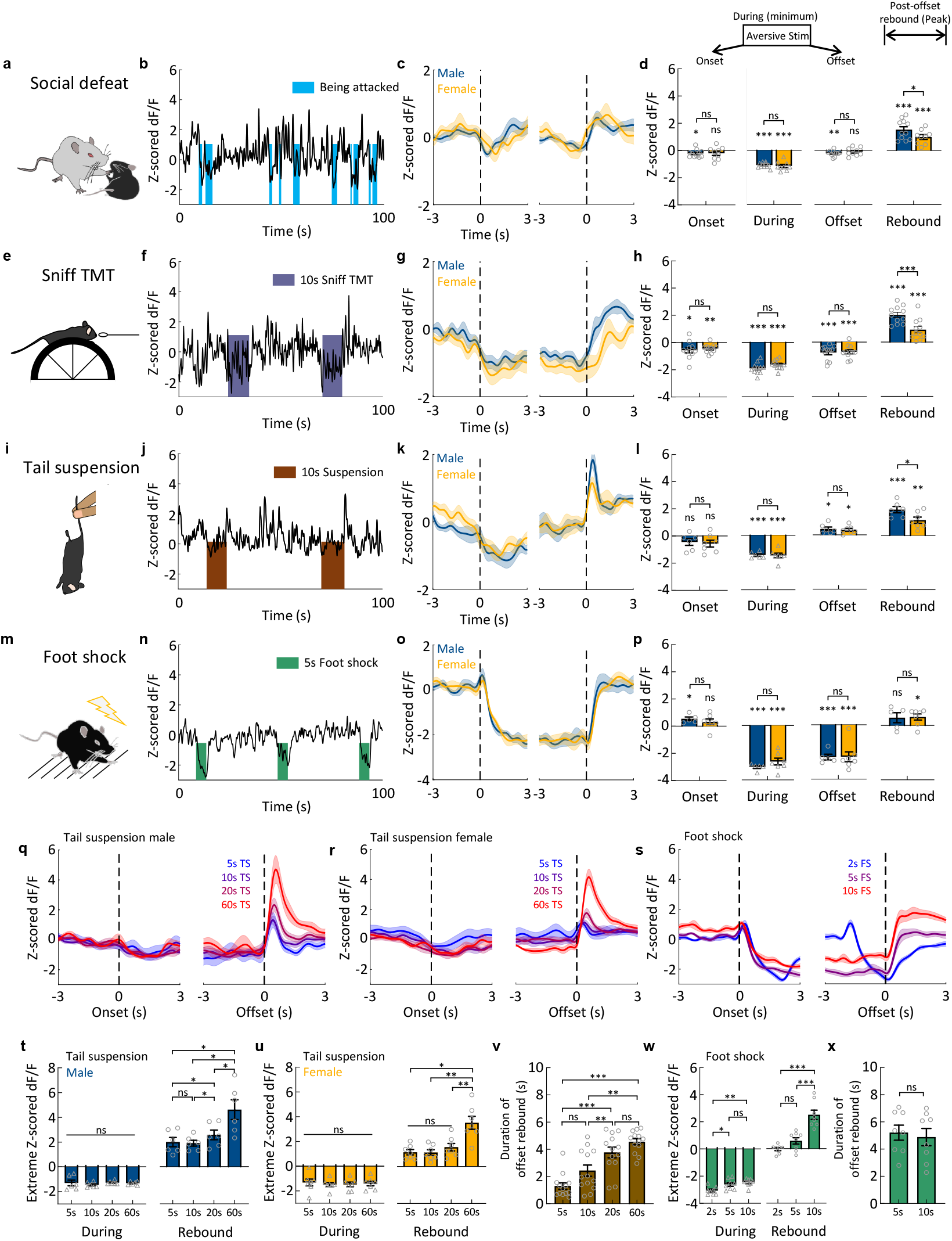
DA responses during aversive social and non-social experiences. **a**, A cartoon illustration of social defeat. **b**, Representative traces of Z scored ΔF/F of GRAB_DA2h_ during defeat. Similar results were observed for 11 male mice and 9 female mice. **c**, Average post-event histograms aligned to the onset (left) and offset (right) of being attacked. Shaded area: s.e.m. (n=11 male mice and n=9 female mice.) **d**, Top: Schematics showing the time periods used for characterizing DA responses related to aversive experiences. Bottom: Summary of Z scored GRAB_DA2h_ responses at the indicated moments of being attacked. Mean ± s.e.m overlaid with individual data points for each group. (n=11 male mice and n=9 female mice. One sample t test with FDR correction revealed significant responses, *p<0.05, **p<0.01, ***p<0.001. Bonferroni’s multiple comparisons following two-way ANOVA revealed the difference in rebound responses between males and females, *p<0.05.) **e**, A cartoon illustration of TMT exposure. **f**, Representative traces of Z scored ΔF/F of GRAB_DA2h_ during sniffing TMT. Similar results were observed for 11 male mice and 11 female mice. **g**, Average post-event histograms aligned to the onset (left) and offset (right) of TMT exposure. Shaded area: s.e.m. (n=11 male mice and n=11 female mice.) **h**, Summary of Z scored GRAB_DA2h_ responses at the indicated moments of sniffing TMT. Mean ± s.e.m overlaid with individual data points for each group. (n=11 male mice and n=11 female mice. One sample t test with FDR correction revealed significant responses, *p<0.05, **p<0.01, ***p<0.001. Bonferroni’s multiple comparisons following two-way ANOVA revealed the difference in rebound response between males and females, ***p<0.001.) **i**, A cartoon illustration of tail suspension. **j**, Representative traces of z-scored ΔF/F of GRAB_DA2h_ during the tail suspension. Similar results were observed from 6 male mice and 7 female mice. **k**, Average post-event histograms aligned to the onset (left) and offset (right) of tail suspension for the tested male and female mice. Shaded area: s.e.m. (n=6 male mice and n=7 female mice.) **l** Summary of Z scored GRAB_DA2h_ responses at the indicated moments of tail suspension. Mean ± s.e.m overlaid with individual data points for each group. (n=6 male mice and n=7 female mice. One sample t test with FDR correction revealed significant responses,*p<0.05, **p<0.01, ***p<0.001. Bonferroni’s multiple comparisons following two-way ANOVA revealed the difference in rebound response between males and females, *p<0.05) **m**, A cartoon illustration of foot shock. **n**, Representative traces of Z scored ΔF/F of GRAB_DA2h_ during 5s foot shock. Similar results were observed from 5 male mice and 7 female mice. **o**, Average post-event histograms aligned to the onset (left) and offset (right) of the 5s foot shock. Shaded area: s.e.m. (n=5 male mice and n=7 female mice.) **p**, Summary of Z scored GRAB_DA2h_ responses at the indicated moments of foot shock. Mean ± s.e.m overlaid for each group. (n=5 male mice and n=7 female mice. One sample t test with FDR correction revealed significant responses, *p<0.05, ***p<0.001. Bonferroni’s multiple comparisons following two-way ANOVA revealed no difference in responses between males and females.) **q-r**, Average post-event histograms aligned to the onset (left) and offset (right) of the tail suspension with different suspension duration in males (**q**) and females (**r**). Shaded area: s.e.m. (n=6 male mice and n=7 female mice.) **s**, Average post-event histograms aligned to the onset (left) and offset (right) of the foot shock with different shock durations. Shaded area: s.e.m. (n=5 males and 3 females.) **t-u**, Maximum GRAB_DA2h_ decrease during tail suspension and maximum post-suspension rebound responses at different suspension durations in males **(t)** and females **(u)**. Mean ± s.e.m overlaid with individual data points for each group. (n=6 male mice: Friedman test for the during period, p=0.43; Tukey’s multiple comparisons following one-way ANOVA revealed the differences in rebound responses between groups, *p<0.05. n=7 female mice: Tukey’s multiple comparisons following one-way ANOVA revealed the differences in rebound responses between groups, *p<0.05, **p<0.01.) **v**, Post-suspension rebound durations vary with the length of tail suspension. Mean ± s.e.m overlaid with individual data points for each group. (n=6 males and 7 females, Tukey’s multiple comparisons following one-way ANOVA revealed the differences in offset rebound duration between groups, **p<0.01, **p<0.001.) **w**, Maximum GRAB_DA2h_ decrease during foot-shock and maximum post-shock rebound responses at different shock durations. Mean ± s.e.m overlaid with individual data points for each group. (n=5 male mice and 3 female mice, Tukey’s multiple comparisons following one-way ANOVA revealed the differences in responses between groups, *p<0.05, **p<0.01, ***p<0.001). **x**, The post-shock rebound duration is similar between 5s and 10s shocks. Mean ± s.e.m overlaid with individual data points for each group. (n=5 male mice and 3 female mice, paired t-test, p=0.69.)

To induce defeat in females, we introduced each recording female mouse to the home cage of a lactating SW female. Similar to that in males, dopamine signal decreased during defeat and rebounded after each defeat episode (**Figure 8b-d**).

We next examined dopamine response during aversive experience that is not socially relevant. We presented 0.05% TMT, a pungent chemical found in fox feces, on a Q-tip to the recorded mouse for 10 minutes when the test animal was head-fixed and positioned on a running wheel (**Figure 8e**); we suspended the test mouse by lifting its tail for 10 seconds to simulate the situation when it was caught by a predator (**Figure 8i**); we also delivered a series of 5-second 0.4 mA foot shocks to elicit pain (**Figure 8m**). In all cases, we observed a consistent decrease in dopamine signal during the aversive experience (**Figures 8e-8p**). For defeat, tail suspension and TMT exposure, the dopamine typically decreased to its lowest level within 2s after the onset and then gradually increases despite the ongoing aversive experience (**Figures 8a-8l**). For foot shock, the dopamine decreased sharply immediately after shock onset and remained maximally suppressed throughout the shocking period (**Figures 8n-8p**). At the offset of all aversive experience, dopamine showed a rebound increase (**Figures 8a-p**). While the decrease in dopamine response during aversive experience was comparable between males and females, the male showed a higher rebound after all aversive experiences except foot shock (**Figures 8d, 8h, 8l and 8p**). Neither dopamine decrease during aversive experience nor the rebound response after the experience showed any adaptation over repeated trials (**Figure 8– figure supplement 1**).

The dopamine rebound may represent a rewarding signal related to relief from aversive experience. We wondered whether the extent of relief (i.e., dopamine rebound), correlates with the negativity of the experience. We thus varied the duration of tail suspension and foot shock and asked whether dopamine responses varied with the length of the aversive experience. While the magnitude of dopamine decrease during tail suspension was similar across all durations, the rebound increase was significantly higher and lasted longer after 60-s suspension than that of shorter durations in both males and females (**Figure 8q-8r and 8t-8v**). Similarly, 10-s foot shock induced significantly higher dopamine rebound than 5-s shock while 2-s shock induced no appreciable rebound, supporting the idea that rebound increase was correlated with the extent of negativity of the experience (**Figure 8s and 8w-8x**).

Lastly, changes in our fluorescence signal could not be accounted by the movement artifacts. In our control animals expressing GFP, we observed no significant change in fluorescence during any behaviors (**Figure 8 – figure supplement 2**).

## Discussion

Using a recently developed genetically encoded dopamine sensor, our study revealed details of dopamine responses during social behaviors and identified sexual dimorphism in the response pattern. We found that dopamine increases in all three phases of social behaviors: approach, investigation and consummation.

The dopamine increase during approach likely signals the motivational state of the animal towards the target. For example, males, not females, show a higher dopamine increase during approaching opposite-sex than same-sex conspecifics, consistent with the fact that males prefer females over males in three-chamber social preference test while females show no preference to either social target (Yao et al., 2017). For another example, mothers and fathers are known to be highly motivated for pups as shown by their willingness to work hard to gain access to pups or giving up other high-value rewards for pups (Hauser and Gandelman, 1985; Lee et al., 2000; Mattson et al., 2001; Wang et al., 2012; Wilsoncroft, 1968). In parents, dopamine increases in NAc core during pup approach is consistently higher than that in non-parental animals. This result is in contrast to the previous finding showing that dopamine responses to pups are similar in males with low and high parental motivation (Lei et al., 2017). This discrepancy is likely due to the low temporal resolution of microdialysis. Although we found that the response during approach differs in fathers and naïve males, dopamine increase during investigation is similar. During infanticide, dopamine level also increased. Thus, the difference in dopamine release during pup approach may be averaged out using microdialysis that has a temporal resolution of 10-20 minutes.

The motivation-dependent release of dopamine during approach is in line with previous functional studies that demonstrated an indispensable role of dopamine in social approach, or more broadly speaking, acquiring access to a social target. For example, blocking the dopamine signaling in the NAc reduces males’ willingness to lever press to gain access to a weaker intruder and attack (Golden et al., 2019a). Increase dopamine release in NAc by activating inputs from the medial preoptic nucleus to the ventral tegmental area promotes pup and male approach in female mice (Fang et al., 2018; McHenry et al., 2017) . Importantly, we found that dopamine rises after the onset of approach in all cases, suggesting that it is not required for approach initiation but might play a role in invigorating the process.

Dopamine also increases during social investigation. This increase is likely driven by the sensory cues from the conspecific and adapts quickly. Similar increase and fast adaption occur during investigation of a non-social target. This response pattern may explain why mice are less interested in familiar conspecific or object than novel ones (Leger et al., 2013; Moy et al., 2004). A previous study suggested that dopamine neurons in VTA show an increase in Ca^2+^ activity as animals withdraw from a novel object (Gunaydin et al., 2014). Here, we found dopamine increases occur mainly during object approach and reaches maximum during object investigation. Although dopamine levels remain elevated above the pre-investigation level at the offset of object investigation, no acute upward change was observed when the animal retreated from the object. This difference in response pattern may be due to discrepancies of dopamine neuron activity and dopamine release at the terminals (Gunaydin et al., 2014).

Dopamine consistently and transiently increases during consummatory phase of social behaviors with little adaption over repeated trials. In general, dopamine rises, reaches peak levels soon after behavior onset, and gradually drops afterwards. At behavior offset, dopamine levels have often returned to baseline, suggesting that dopamine signals may be most important for marking transitions in behaviors instead of their maintenance. The sex dimorphism of dopamine release is most noticeable during the consummatory phase. During attack, males show higher dopamine increase than postpartum female mice. While dopamine increases during all phases of sexual behaviors in males, only receptive, not unreceptive, females show dopamine increase and only when she is initially mounted and soon after male ejaculation. The sex difference in dopamine release during aggression and sexual behaviors likely stem from the sex differences in behaviors themselves. Indeed, aggressive behaviors in male and female mice are triggered by different sensory cues while sexual behaviors differ between sexes in both sensory triggers and motor execution. Regardless of the cause of sex differences in dopamine release, this difference in release pattern likely signals different valence of the experience and reinforces the behavior in a sex specific way. Indeed, while in male mice repeated attacks lead to an increase in aggression, a phenomenon known as winner effect, female mice have not been found to show a clear winner effect (Hashikawa et al., 2018).

What is the function of dopamine release during the consummatory social behaviors? One likely role of dopamine is for reward associative learning. Inputs to NAc cells that carry information regarding the environmental contexts and mating partners could be strengthened by the dopamine release during consummatory social actions and in turn facilitate approach to the cues and enhance the chance to be engaged in similar social behaviors in the future (Aragona et al., 2003; Aragona et al., 2006; Balfour et al., 2004; Gingrich et al., 2000; Meisel et al., 1996). Beyond reward learning, does dopamine release also play a role in the expression of consummatory social behaviors? Answer to this question could be behavior-specific. Pup retrieval, for example, has been shown to be critically dependent on dopamine receptor activation in the NAc shell (Keer and Stern, 1999; Numan et al., 2005). Blocking D1 receptor signaling in NAc disrupts pup retrieval whereas D1 receptor agonist in the NAc facilitates the onset of maternal behavior (Numan et al., 2005; Stolzenberg et al., 2007). Inhibiting D1R expressing cells in NAc reduced attack duration in male mice (Golden et al., 2019b). In contrast, male sexual behaviors are not affected by dopamine depletion in the NAc although manipulated males showed a decrease in noncontact erection, suggesting a decrease in sexual motivation (Liu et al., 1998; Moses et al., 1995). The differential importance of dopamine signaling in the expression of various social behaviors suggests distinct roles for the NAc in driving these behaviors. Indeed, NAc has been suggested as a key part of the maternal circuit but largely left out of consummatory sexual behavior circuits (Jennings and de Lecea, 2020; Kohl et al., 2017; Numan, 2007). Of note, deficits in social reward conditioning are mainly caused by D2 receptor antagonists in the NAc while deficits in social behavior expression mainly resulted from D1 receptor antagonism (Aragona et al., 2006; Gingrich et al., 2000; Golden et al., 2019b; Meisel et al., 1996; Numan et al., 2005). Thus, it is possible that dopamine release in the NAc serves dual roles: it promotes the execution of certain social behaviors through D1R and facilitates reward conditioning through D2R.

Previous microdialysis study showed an overall increase in dopamine during defeat (Tidey and Miczek, 1996). Here, using a method with finer temporal resolution, we observed a decrease in extracellular dopamine during defeat followed by rebound increase after the defeat terminates. Indeed, dopamine rebound is commonly observed at the termination of an aversive experience regardless of its exact nature and the magnitude of rebound is positively correlated with the duration of the experience. This suppression-rebound response pattern is in line with previous electrophysiological recordings in the VTA and FSCV recording in the NAc (Brischoux et al., 2009; Budygin et al., 2012). Behavioral experiments also demonstrated that relief from the pain can be used as a “reward” to induce preference to the associated context or sensory cues (Navratilova et al., 2015). We speculate the dopamine rebound at the end of defeat could contribute to the rapid emergence of avoidant behavior towards the aggressor after a short period of defeat as the rebound often occurs when the defeated animal moves away from the aggressor. Interestingly, we did not observe a rebound increase at the termination of forced mounting in non-receptive females and females do not develop avoidance towards the male afterwards. In addition to the rebound increase after aversive experience, we also noticed a transient suppression in dopamine activity after the termination of positive social behavior, such as mounting and intromission in males and pup retrieval in fathers. The biphasic activity change at the onset and offset of an experience may together facilitate learning of cues predictive of the transition of behaviors.

In short, our study demonstrated fast and dynamic dopamine responses in the NAc during various phases of social behaviors. The functional role of dopamine is likely to be complex, multifaceted and across both fast and slow time scales. Future studies with temporally precise manipulation tools that can block and enhance the dopamine release in a behavior-locked manner will facilitate our understanding of the mesolimbic dopamine function in social behaviors.

## Material and Methods

### Animals

All procedures were approved by the IACUC of NYULH in compliance with the NIH guidelines for the care and use of laboratory animals. Mice were housed at 18–23 °C with 40–60% humidity under a 12 h light–dark cycle (dark cycle, 10 p.m. to 10 a.m.), with food and water available ad libitum. Test animals were adult Drd1-cre (> 8 weeks, MMRRC_030989-UCD). Ai9 mice (Jackson stock no.007909) were crossed with Drd1-cre mice for revealing D1R expression. Stimulus animals were adult C57BL/6N male and female mice, adult BALB/c male and female mice, or sexually experienced C57BL/6N and Swiss Webster males (aggressor) purchased from Charles River, and 3-7 days old pups were bred from test mice or wildtype C57BL/6N breeders. After surgery, all the animals are single housed. All experiments were performed during the dark cycle of the animals.

### Immunofluorescence

For histological analysis, animals were deeply anesthetized and transcardially perfused with 20 mL of PBS, followed by 20 mL of 4% paraformaldehyde (Electron Microscopy Sciences, cat. no. 15714) in PBS. After perfusion, brains were harvested, post-fixed in 4% paraformaldehyde for 4h at 4 °C and cryoprotected in 20% (w/v) sucrose for 24h. The brains were then embedded in an O.C.T compound (Fisher Healthcare, cat. no. 23730571) and sectioned into 60-μm-thick slices using a CM1900 cryostat (Leica). GRAB_DA2h_ was immunostained using a chicken anti-GFP antibody (1:1,000, Abcam, cat. no. ab13970) followed by an Alexa 488-conjugated donkey anti-chicken secondary antibody (1:1,000, Jackson ImmunoResearch, cat. no. 116967). DAPI (1:20,000; Thermo Fisher, cat. no. D1306) was used with the secondary antibody to visualize the nucleus. The GRAB_DA2h_ fluorescence images were acquired with a virtual slide microscope (Olympus, VS120) in 10x mode or a confocal microscope (Zeiss LSM 510 or 700 microscope) for the high-resolution image in 40x mode.

### Fiber photometry

The male mice were screened for aggression before the surgery. For the test mouse with a single fiber implanted, 80 nL AAV9.hSyn.DIO. GRAB_DA2h_ (Vigene Biosciences, titer: 4.97e+13 gc/ml) was injected into one side of NAc core (anterior-posterior (AP): +0.98 mm relative to Bregma; medial-lateral (ML): ±1.2mm relative to Bregma; dorsal-ventral (DV): 4.2 mm from brain surface) using a nanoinjector (World Precision Instruments, Nanoliter 2000). For bilateral recording, AAV9.hSyn.DIO. GRAB_DA2h_ (Vigene Biosciences) was injected into one side and AAV9.hSyn.DIO. GRAB_DA2h_ or AAV9.hSyn.loxP. GRAB_DA2h_.loxP (Vigene Biosciences, titer: 1.43e+13 gc/ml) was injected into the contralateral side of NAc core for the D1-D1 mice or D1-nonD1 mice respectively. Control mice were injected with AAV2.CAG.Flex.GFP (UNC, titer: 4.00e+12 gc/ml) bilaterally. After virus injection, a custom-made optic fiber assembly (Thorlabs, FT400EMT and SFLC440-10) was implanted approximately 300 μm above each injection site. Fiber photometry recording was performed two weeks after AAV injection. The setup used for recording was constructed as described previously (Falkner et al., 2016). In brief, a 400-Hz 472-nm bandpass (passing band: 472 ± 15 nm, FF02-472/30-25, Semrock) filtered light-emitting diode (Thorlabs, LED light: M470F1; LED driver: LEDD1B) was used to excite GRAB_DA2h_ or GFP. The emission light collected from the recording site was bandpass filtered (passing bands: 535 ± 25 nm, FF01-535/505, Semrock), detected by a Femtowatt Silicon Photoreceiver (Newport, 2151), and recorded using a real-time processor (RZ5, TDT). The 400-Hz signals carrying fluorescence intensity of GRAB_DA2h_ or GFP were extracted in real time using a custom TDT program. To analyze the recording data, the MATLAB function ‘‘msbackadj’’ with a moving window of 25% of the total recording duration was first applied to obtain the instantaneous baseline signal. The instantaneous ΔF/F was calculated as (F_raw_ –F_baseline_)/F_baseline_. The Z scored ΔF/F of the entire recording session was calculated as (ΔF/F – mean(ΔF/F))/std(ΔF/F). The peri-event histogram (PETH) of a given behavior was plotted by aligning the Z scored ΔF/F signal to the onset or offset of the behavior. The response during each behavior episode is defined as the maximum Z during the behavior if the average Z increases during the behavior and the minimum Z if average Z decreases during the behavior. The onset and offset response are determined as the Z values at the moment of behavior onset and offset. The post-offset response is defined as the average Z between 0-1 second after the end of the behavior. The post-offset rebound response is the maximum Z 0-2 seconds after the termination of aversive experience. The velocity of the animal was calculated as the displacement of the body location of the animal between every other frame, and the acceleration was calculated as the velocity difference between two adjacent frames. The onset of the movement is defined as the movement following at least 2s immobility (velocity < 2 cm/s) and reaches a velocity of minimally 15 cm/s for at least 1.5 s. The correlation coefficient was calculated using MATLAB function ‘corrcoef’.

### Behavioral paradigm and analysis

Animal behaviors in all experiments were video recorded from both the side and top of the cage using two synchronized cameras (Basler, acA640-100 gm) and a commercial video acquisition software (StreamPix 8, Norpix) in a semi-dark room with infrared illumination at a frame rate of 25 frames/s. Manual behavioral annotation was performed on a frame-by-frame basis using custom software written in MATLAB (https://pdollar.github.io/toolbox/). DeepLabCut was used for animal tracking (Mathis et al., 2018)

#### Social interaction between animals of the same sex

An adult BALB/c male was introduced to the home cage of the test male mouse, or an adult BALB/c female was introduced to the home cage of the test female mouse. If the test female is lactating, pups were removed 10 minutes prior to the intruder introduction. During social encounters, we identified three behaviors of the test mice --approach, investigation and attack. “Approach” was defined as continuous movement toward a stationary intruder mouse until the center mass of the two animals are below 100 pixels. “Investigation” was defined as close contact to any part of the intruder’s body. ‘‘Attack’’ was defined as a suite of intense actions aiming at biting the intruders, including pushes, lunges, bites, tumbling, and fast locomotion episodes between such movements.

#### Social interaction between animals of different sexes

An adult receptive or unreceptive female intruder mouse was introduced into the home cage of the test male mouse. For the test female mouse, an adult sexually experienced male mouse was introduced into the female’s home cage. During social encounters, we annotated “approach” and “investigation” of the test mouse, and “mount”, “intromit” and “ejaculate” of the male mouse. “Mount” is when the male grasped and mounted the female’s flanks. “Intromit” includes both rapid thrust against the female’s rear and deep rhythmic thrust. “Ejaculate” starts when the male suddenly ceases all thrusting movements but still holding onto the female’s flank and then after a few seconds slumps to the side of the female. “Ejaculate” ends when the male resumes movements.

#### Food Intake

A cup of peanut butter (around 2 grams, Jif Creamy Peanut Butter) was introduced into the home cage of the test mouse. The test mouse freely interacted with the peanut butter for 10 minutes. During the interaction, we identified “approach” and “eat”. “Approach” is defined as continuous movement toward the peanut butter cup until the mouse head is above the cup. “Eat” is defined as active licking and chewing peanut butter.

#### Pup-related behaviors

5 -7 pups from test mice or C57BL/6N breeders were introduced, one at a time, into the home cage of the test mouse for a total of 10 minutes. Approach, close interaction, biting, and retrieval are annotated. “Approach” is defined as continuous movement toward a pup until the head of the test mouse is above the pup. “Close interaction” is defined as close contact with pups including sniffing, licking, and grooming. “Biting” is when the test mouse holds and bites the pup and causes harm. “Retrieval” starts when the mouse picks up a pup with its mouth and ends when it drops the pup into or around the nest.

#### Social defeat

To induce defeat, the male test mouse was introduced into a Swiss Webster male’s (aggressor) cage for 10-15 minutes. The female test mouse was introduced into a lactating Swiss Webster female’s cage. During interaction with the aggressor, we annotated “being attacked” of test mice, which is defined as when the aggressor attacks the test mouse. No other behavior tests were performed after social defeat for at least 24 hours.

#### Footshock

The test mouse was placed in a fear conditioning chamber (Med Associates). After 5 minutes habituation, a series of electric shocks were delivered through the floor grids (2-s shock: 0.4 mA for 2 s, 40 s interval, 6 times; 5-s shock: 0.4 mA for 5 s, 60 s interval, 4 times; 10-s shock: 0.4 mA for 10 s, 90 s interval, 4 times). The shock was controlled by TTL pulses generated using a real-time processor (RZ5, TDT). No other behavior tests were performed after footshock for at least 24 hours.

#### TMT exposure

The test mouse was habituated on a 3D printed head-fixed apparatus 10 minutes a day for 3 days. On the test day, 0.05% TMT (diluted in mineral oil) was delivered to the head-fixed mouse on a Q-tip moving along a linear track for 6 times. Each TMT representation lasted for 10 s with a 120-s between trials. The onset of TMT exposure was defined as when the Q-tip reached the closest point (< 1cm) to the mouse nostrils. The offset was defined as when the Q-tip started to retract. No other behavior tests were performed after TMT exposure for at least 24 hours.

#### Tail suspension

During the test, the experimenter grabbed the tail of the mouse and lifted it approximately 25 cm above the floor of the mouse cage. 5-s tail suspension was first performed for 6 times with 120-s between trials. Then, 10-s, 20-s, 60-s tail suspension trials were performed with 180 s between trials. No other behavior tests were performed after tail suspension for at least 24 hours.

### Drug Injection

The test mouse was habituated for head fixation for 3 continuous days. On the test day, the mouse was head-fixed while DA signals is continuously monitored. After 10 minutes baseline, we intraperitoneally injected 0.9% NaCl (10ml/kg, Hanna Pharmaceutical Supply, cat. no. NC9054335) and 40 minutes later, 20 mg/kg GBR 12909 (Tocris, cat. no. 0421) was injected and the signal was recorded for at least 30 minutes after injection.

### Statistics

All the data were tested for normality first by using Kolmogorov-Smirnov test. If all the data was normally distributed, paired t tests were performed for comparisons between two groups within the animal, unpaired t tests between animals, one-way ANOVA with Turkey’s post hoc test, or ordinary (or mixed effect) two-way ANOVA with Bonferroni’s multiple-comparison post hoc test were performed for comparisons between groups. In addition, one sample t-test was performed to determine whether the Z score is significantly different from 0, followed by false discovery rate correction with a false discovery rate of 0.05. If one or more groups were not normally distributed, Mann–Whitney test, Wilcoxon matched-pairs signed-rank test, the Friedman test with Dunn’s multiple-comparison post hoc test, or repeated-measures two-way ANOVA with Bonferroni’s multiple-comparison test were performed. Wilcoxon rank test was used to compare the means of two groups or the mean of a single group with 0, followed by false discovery rate correction with a false discovery rate of 0.05. Details of each statistical test can be found in the **supplementary table 1**.All error bars or error shades represent ± SEM. *, p < 0.05; **, p < 0.01; ***, p < 0.001.

## Acknowledgements

We thank Dr. Nic Tritsch for feedbacks on the manuscript, Yiwen Jia for helping with genotyping, Yijing Gao for initial testing of dopamine sensors and Jiawen Fan for assisting with behavior annotation.

## Competing Interests

The authors declare no competing interests.

---

**Figure 1 - figure supplement 1.**
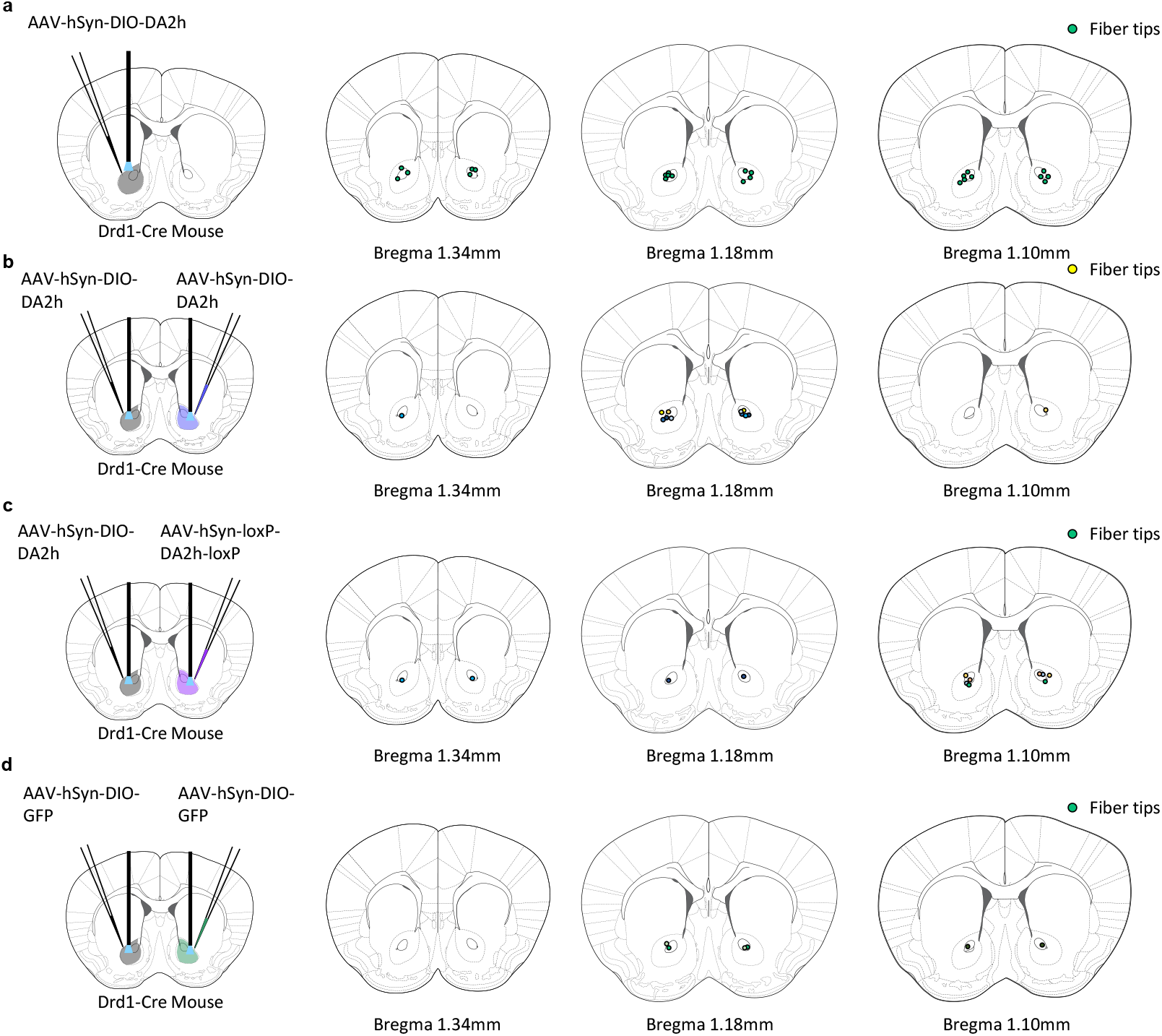
Optic fiber placements for recordings. **a-d**, Experimental design (left) and coronal brain sections (right) at the bregma level of NAc showing the optic fiber ends. Each dot represents one animal in **(a)**, and dots with same color in **(b-d)** represent same animal. Brain atlas images are modified from (Franklin and Paxinos, 2013).

**Figure 2 - figure supplement 1.**
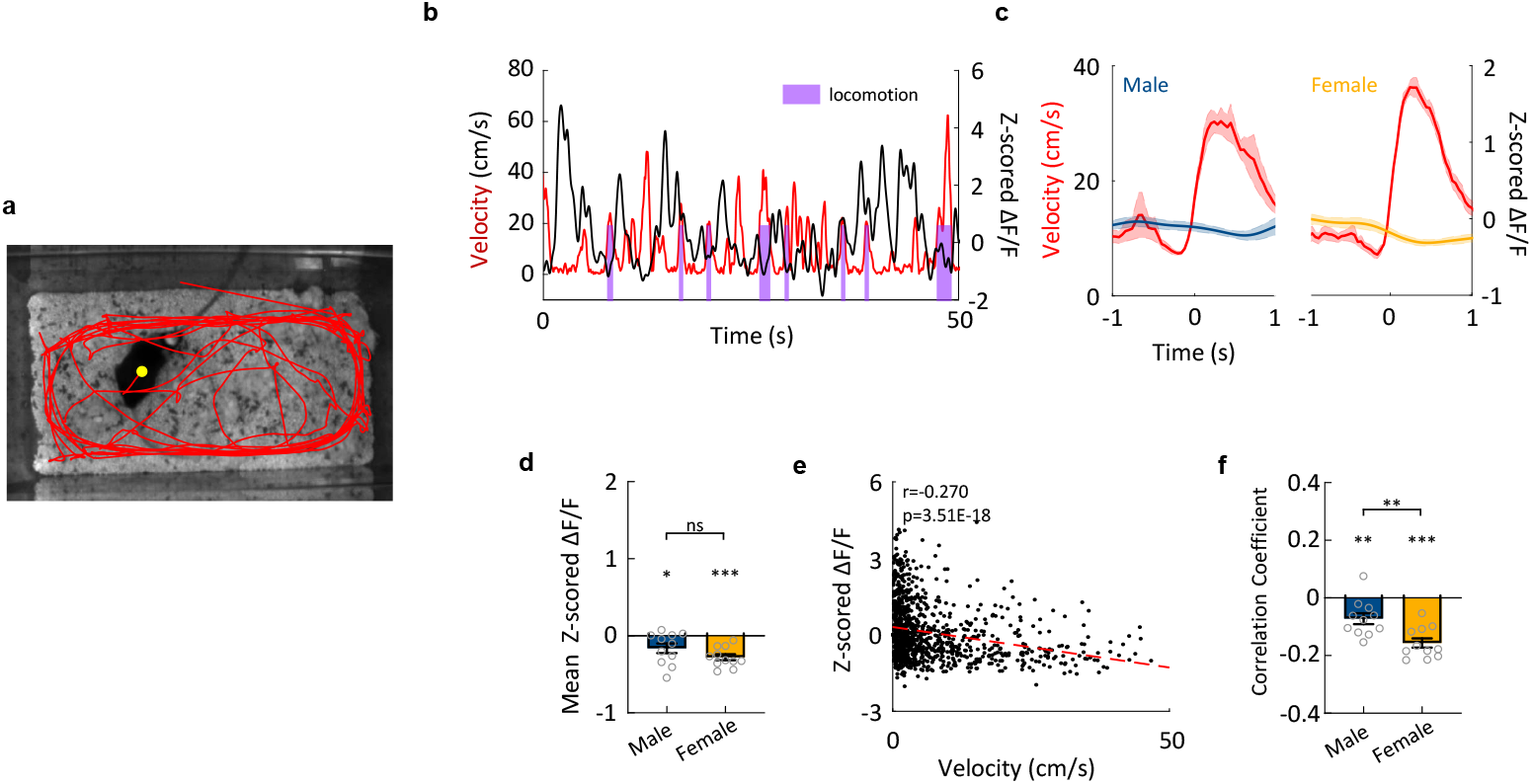
No increase in DA activity at the onset of locomotion or acceleration. **a**, Representative trajectory tracked by DeepLabCut (Mathis et al., 2018). **b**, Representative traces of velocity (red) and z-scored ΔF/F of GRAB_DA2h_ (black) from an example male mouse. Color shades indicate periods of locomotion. Similar results were observed from 11 male mice and 11 female mice. **c**, Average post-event histograms of velocity (red) and GRAB_DA2h_ signal aligned to the locomotion onset of males (left) and females (right). Shaded area: s.e.m. (n=11 male mice and n=11 female mice.) **d**, Averaged Z scored GRAB_DA2h_ signal during 0-1s of all locomotion episodes. (n=11 male mice and n=11 female mice. One sample t test revealed significant activity changes, *p<0.05, ***p<0.001. Unpaired t-test found no difference between two groups, p=0.12.) **e**, A scatter plot showing the correlation between velocity and Z scored ΔF/F from one example female mouse. Similar results were observed from 11 male mice and 11 female mice. (A thousand data points were randomly selected for visualization, Pearson correlation, r = -0.27, p=3.15E-18.) **f**, Summary of correlation coefficient between velocity and Z scored ΔF/F. (n=11 male mice and n=11 female mice. One sample t test revealed the correlation coefficients are significantly negative across animals, **p<0.01, ***p<0.001. Unpaired t-test found a difference between males and females, p = 0.003.)

**Figure 5 - figure supplement 1.**
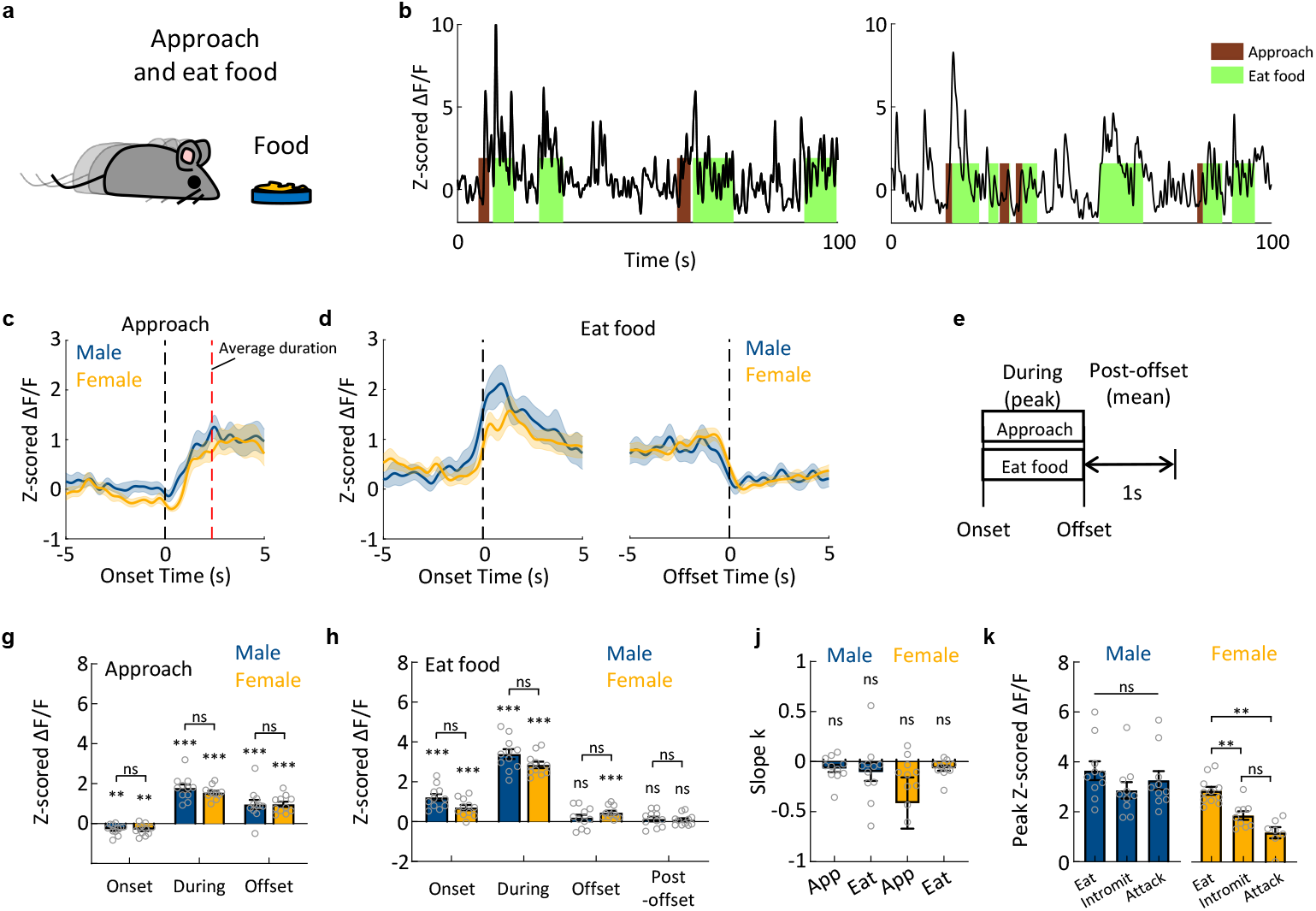
DA responses during food intake. **a**, A cartoon illustration of mouse approaching food. **b**, Representative traces of z-scored ΔF/F of GRAB_DA2h_ during approach and eat peanut butter. Similar results were observed from 11 male and 11 female mice. **c**, Average post-event histograms aligned to the onset of approaching food. Shaded area: s.e.m. (n=11 male mice and n=11 female mice.) **d**, Average post-event histograms aligned to the onset (left) and offset (right) of eating. Shaded area: s.e.m. (n=11 male mice and n=11 female mice.) **e**, Schematics showing the periods used for characterizing DA responses related to food intake. **g**, Summary of Z scored GRAB_DA2h_ responses at the onset and offset of, and during approach food. Mean ± s.e.m overlaid with individual data points for each group. (n=11 male mice and n=11 female mice. One sample t test followed by FDR correction to reveal significant responses, **p<0.01, ***p<0.001. Two-way ANOVA group x time interaction, F(2,40)=0.6226, p=0.54.) **h**, Summary of Z scored GRAB_DA2h_ responses at the onset and offset of, and during and after eating food. (n=11 male mice and n=11 female mice. One sample t test followed by FDR correction to reveal significant responses, ***p<0.001. Bonferroni’s multiple comparisons following two-way ANOVA revealed no significant difference in responses between males and females.) **j**, Summary of slope k of GRAB_DA2h_ responses over repeated eating episodes. (n=11 male mice, n=11 female mice. One sample t test followed by FDR correction to reveal significant adaptations.) **k**, Summary of Z scored GRAB_DA2h_ responses during food intake, intromission, and attack. (n=10 male mice, one-way ANOVA, F(1.340,12.06)=1.166, p=0.32. n=11 female mice for food intake, n=9 female mice for intromission, n=7 female mice for attack, Tukey’s multiple comparisons following one-way ANOVA revealed differences in responses between groups in females, **p<0.01.)

**Figure 5 - figure supplement 2.**
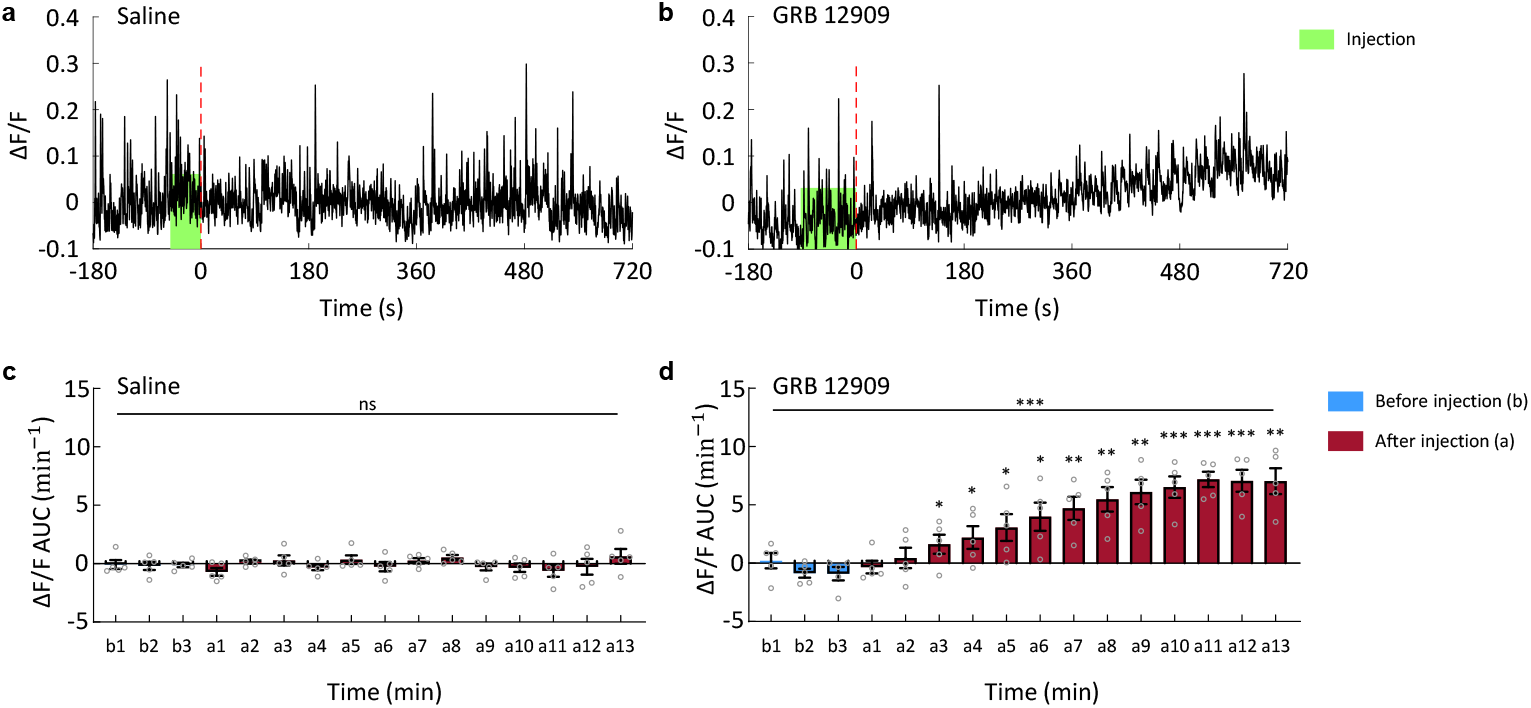
Administration of DAT inhibitor induces a sustained increase in GRAB_DA2h_ signal. **a-b**, Representative traces of z-scored ΔF/F of GRAB_DA2h_ after i.p. injection of saline (left) and 20 mg/kg GBR 12909 (right). Similar results were observed from 5 male mice. **c**, Accumulated DA signals before and after saline treatment. Mean ± s.e.m overlaid with data points from individual animals for each group. (n = 5 mice, Friedman test, p=0.58.) **d**, Accumulated DA signals before and after GBR 12909 treatment. Mean ± s.e.m overlaid for each group. (n = 5 mice, Tukey’s multiple comparisons following one-way ANOVA revealed the difference between b3 and other time periods, *p<0.05, **p<0.01, ***p<0.001.)

**Figure 8 - figure supplement 1.**
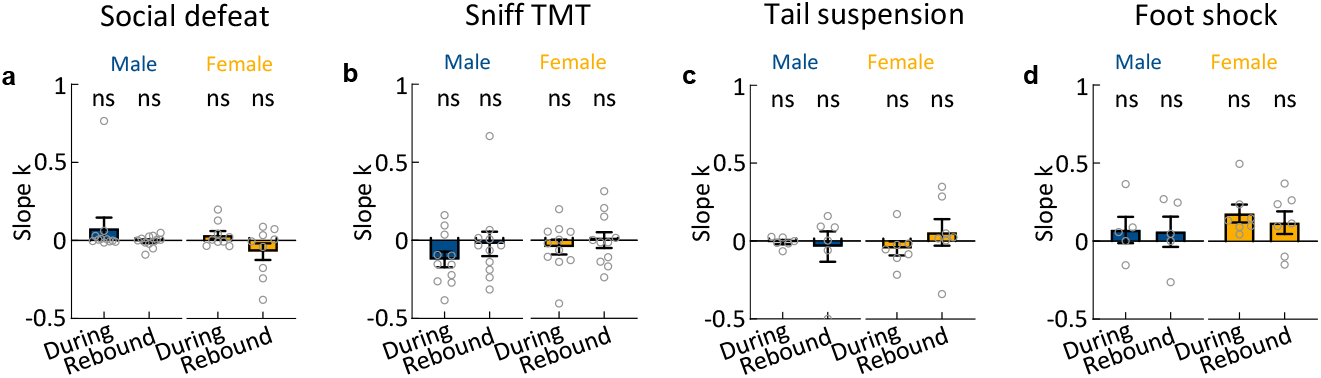
Dopamine responses during and immediately after aversive experiences do not adapt over repeated trials. **a**, Summary of slope k of GRAB_DA2h_ responses over repeated defeat events. Mean ± s.e.m overlaid with individual data points for each group. (n=11 male mice and n=female mice. One sample t test followed by FDR correction revealed no significant adaption.) **b**, Summary of slope k of GRAB_DA2h_ responses over repeated TMT exposure. Mean ± s.e.m overlaid with individual data points for each group. (n=11 male mice and n=11 female mice. One sample t test followed by FDR correction revealed no significant adaption.) **c**, Summary of slope k of GRAB_DA2h_ responses over repeated tail suspension. Mean ± s.e.m overlaid with individual data points for each group. (n=6 male mice and n=7 female mice. One sample t test followed by FDR correction revealed no significant adaption.) **d**, Summary of slope k of GRAB_DA2h_ responses over repeated 5s foot-shock. Mean ± s.e.m overlaid with individual data points for each group. (n=5 male mice and n=7 female mice. One sample t test followed by FDR correction revealed no significant adaption.)

**Figure 8 - figure supplement 2.**
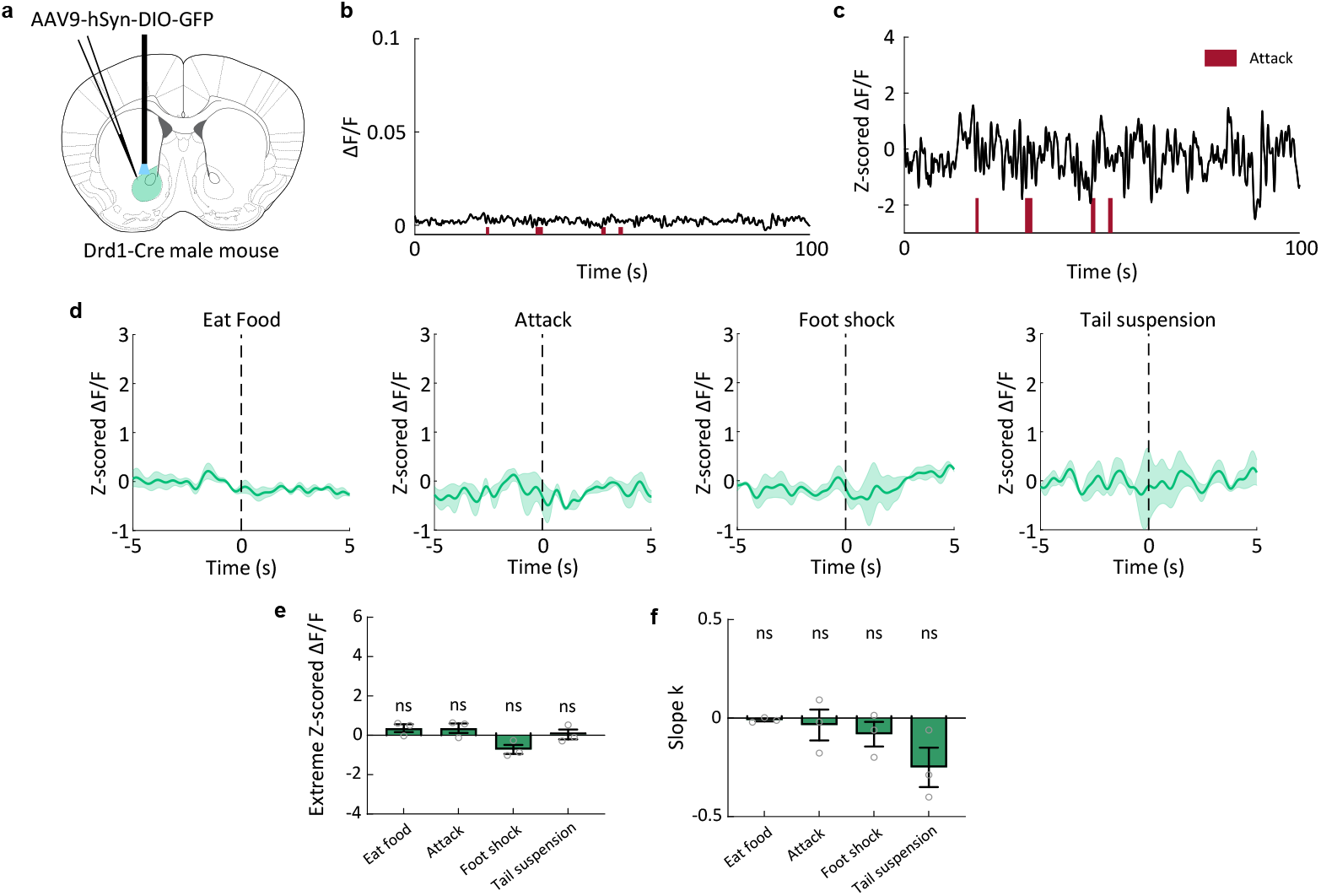
No change in fluorescence signal during any behaviors in GFP control animals. **a**, Schematics illustrating the experimental design. Brain atlas images are modified from (Franklin and Paxinos, 2013). **b-c**, Representative traces of ΔF/F **(b)** and Z scored ΔF/F **(c)** of GFP during inter-male aggression. Similar results were observed for 3 male mice. **d**, Average post-event histograms aligned to the onset of eat, attack, foot-shock, and tail suspension. Shaded area: s.e.m. (n = 3 male mice.) **e**, Maximum GFP increase during food intake and attack, and maximum GFP decrease during tail suspension and foot shock. Mean ± s.e.m overlaid with individual data points for each group. (n = 3 male mice. One sample t test followed by FDR correction revealed no significant response.) **f**, Summary of slope k of GFP responses over repeated events. Mean ± s.e.m overlaid for each group. (n=3 male mice. One sample t test followed by FDR correction revealed no significant adaption.)

**Supplementary Table 1.**
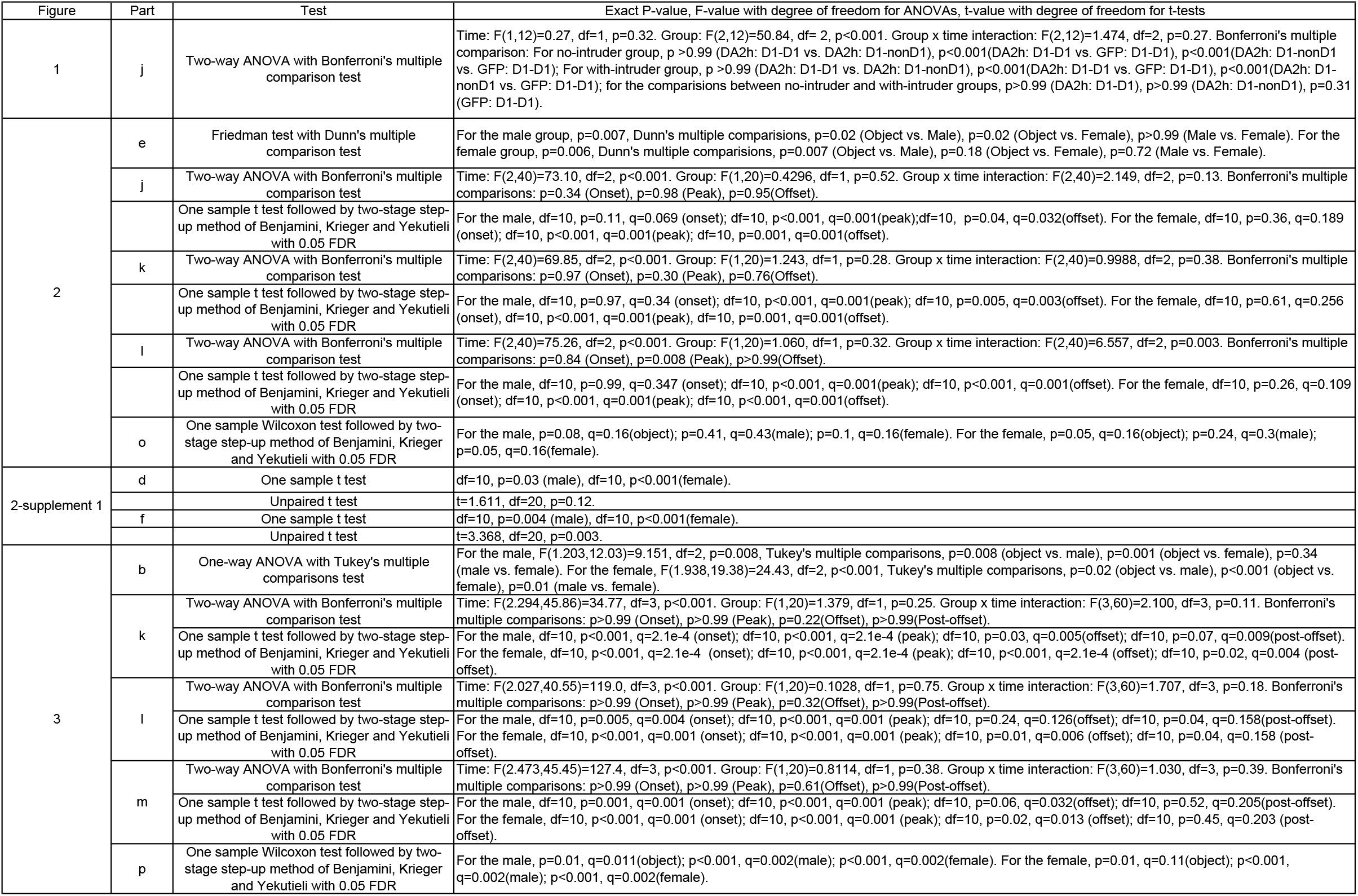
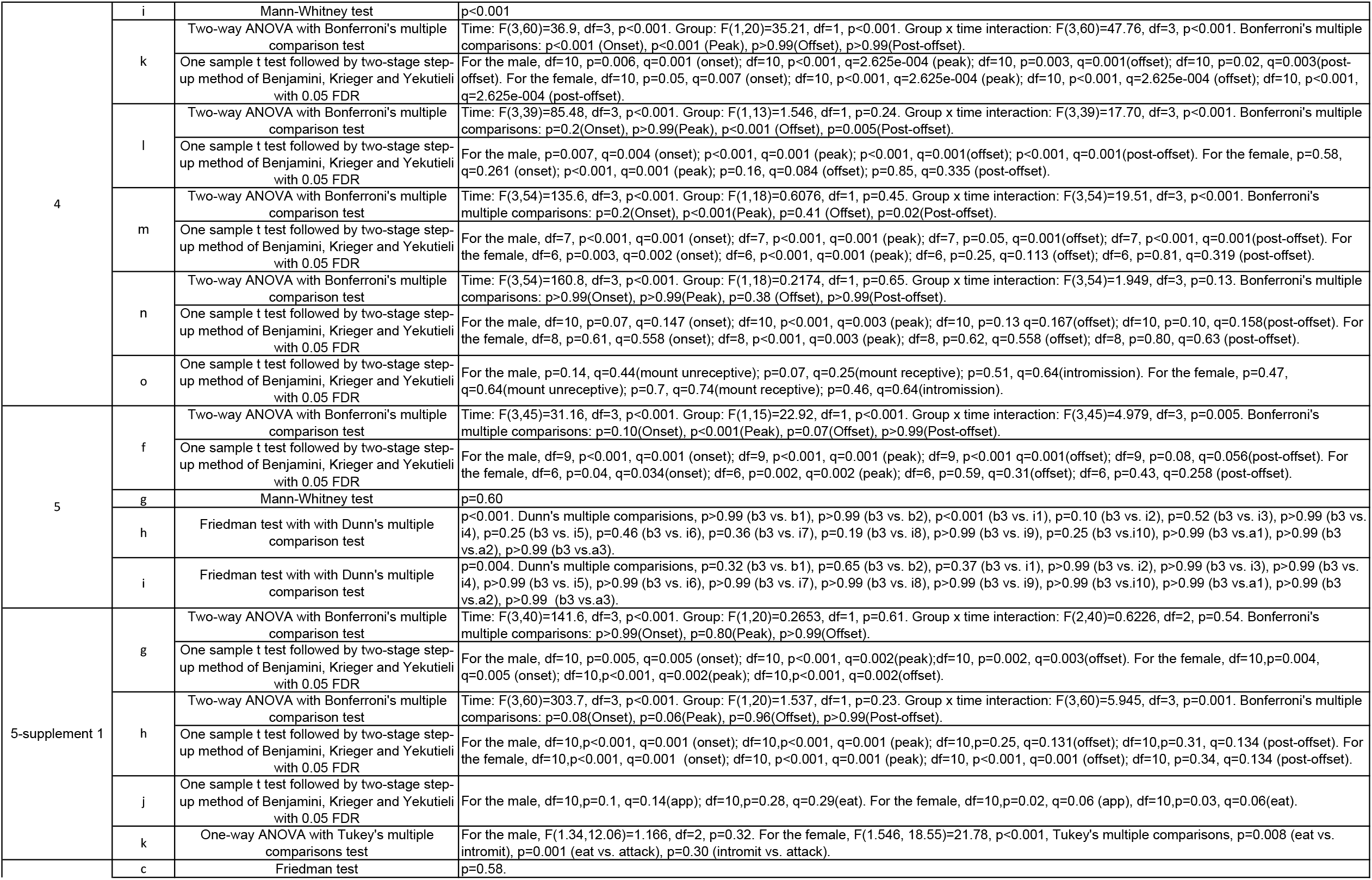
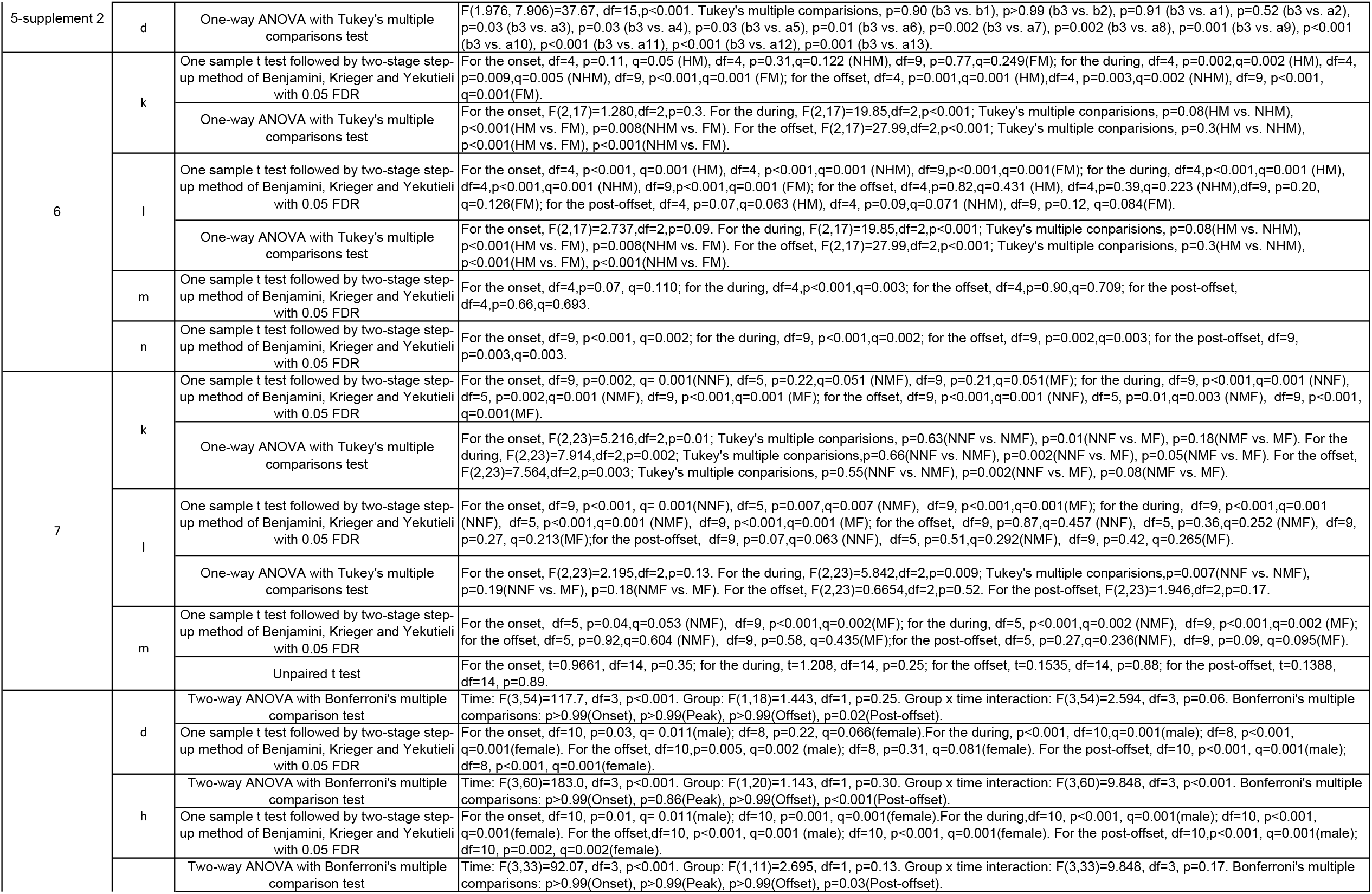
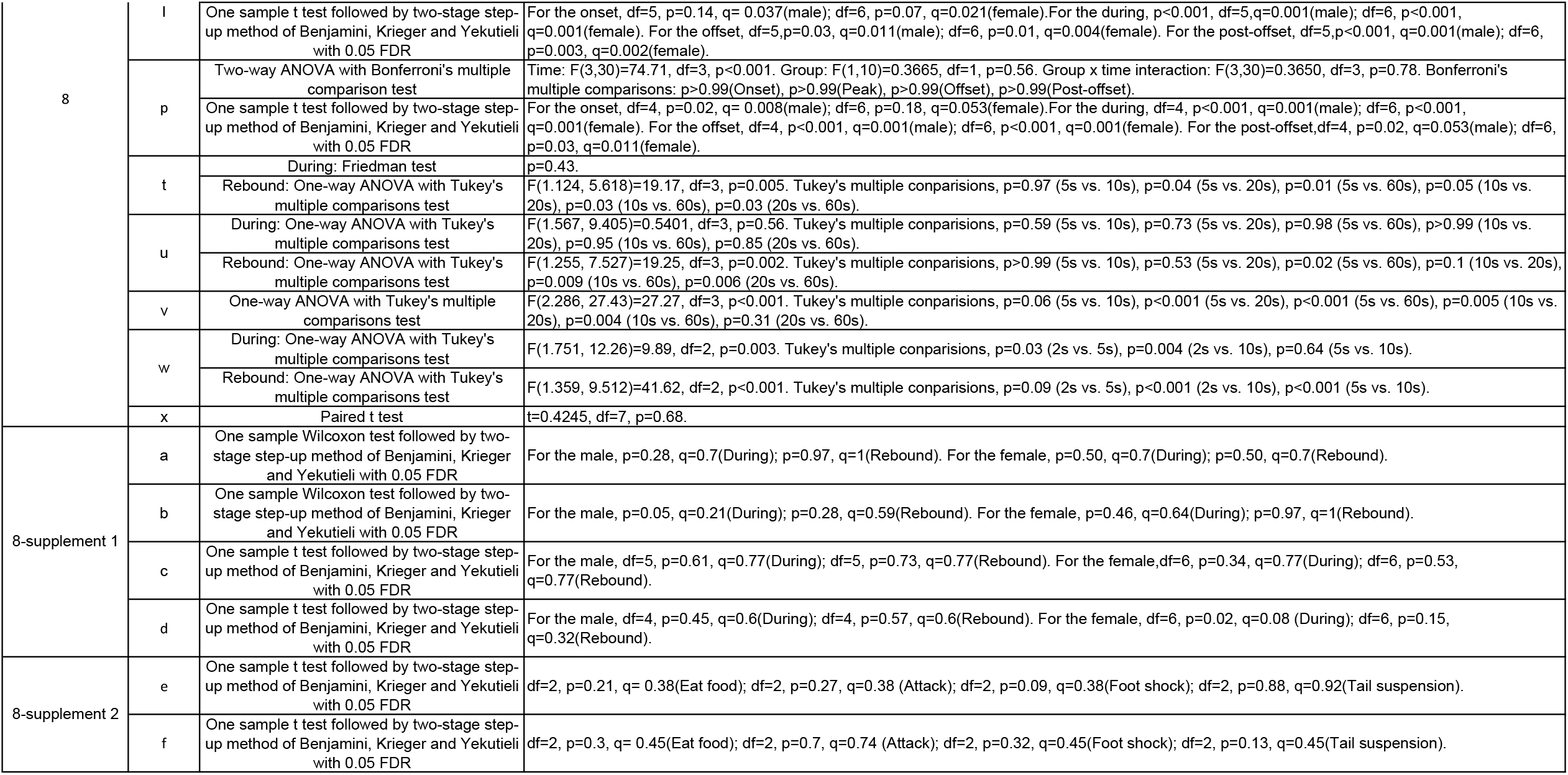

## Notes

### Competing Interest Statement

The authors have declared no competing interest.

